# Bacterial growth under confinement requires transcriptional adaptation to resist metabolite-induced turgor pressure build-up

**DOI:** 10.1101/2024.09.20.614086

**Authors:** Laure Le Blanc, Baptiste Alric, Romain Rollin, Laura Xénard, Laura Ramirez Finn, Sylvie Goussard, Laurent Mazenq, Molly A. Ingersoll, Matthieu Piel, Jean-Yves Tinevez, Morgan Delarue, Guillaume Duménil, Daria Bonazzi

**Affiliations:** Institut Pasteur, Université Paris Cité, INSERM U1225, Paris, France; LAAS-CNRS, Université de Toulouse, CNRS, Toulouse, France; Institut Pierre Gilles de Gennes, Institut Curie, CNRS, Paris, France; Institut Pasteur, Université Paris Cité, Image Analysis Hub, Paris, France; Université Paris Cité, INSERM U1016, CNRS UMR 8104, Institut Cochin, Paris, France; Institut Pasteur, Université Paris Cité, Department of Immunology, Paris, France

**Author notes:** Current address: School of Life Sciences, École Polytechnique Fédérale de Lausanne (EPFL), Lausanne, Switzerland. These authors contributed equally (Baptiste Alric, Romain Rollin, Laura Xénard). These authors jointly supervised this work (Morgan Delarue, Guillaume Duménil, Daria Bonazzi).

**Keywords:** Spatial confinement, Bacterial physiology, Mechanical constraints, Mechanomicrobiology, Microfluidics, Macromolecular crowding, Rcs envelope stress response, *Escherichia coli*, Urinary tract infection

## Abstract

Bacterial proliferation often occurs in confined spaces, during biofilm formation, within host cells, or in specific niches during infection, creating mechanical constraints. We investigated how spatial confinement and growth-induced mechanical pressure affect bacterial physiology. Here, we found that, when proliferating in a confining microfluidic-based device with access to nutrients, *Escherichia coli* cells generate forces in the hundreds of kPa range. This pressure decouples growth and division, producing shorter bacteria with higher protein concentrations. This leads to cytoplasmic crowding, which ultimately arrests division and stalls protein synthesis. In this arrested state, the pressure produced by bacteria keeps increasing. A minimal theoretical model of bacterial growth predicts this novel regime of steady pressure increase in the absence of protein production, that we named *overpressurization*. In this regime, the Rcs pathway is activated and that abnormal shapes appear in *rcs* mutant populations only when they reach the overpressurized state. A uropathogenic strain of *E. coli* displayed the same confined growth phenotypes *in vitro* and requirement for Rcs in a mice model of urinary tract infection, suggesting that these pressurized regimes are relevant to understand the physiopathology of bacterial infections.

## INTRODUCTION

Mechanical forces shape the development and condition of all forms of life. Animal cells probe and apply forces on their environment, and their capacity to respond to mechanical signals, also called mechanosensing, regulates diverse processes such as growth, motility, state of differentiation, and behavior within tissues(*1*). Dysregulation of conserved mechanotransduction pathways is involved in the emergence of several diseases impacting tissue development and homeostasis, such as cancer(*2*, *3*). Despite a major focus of the mechanobiology field on eukaryotic organisms, in recent years mechanical sensing and adaptation to forces proved essential also for microbes, especially among bacteria to colonize diverse ecological niches(*4*). Recent technological advances in live imaging and microfabrication have opened the path to the investigation of how single bacterial cells respond to mechanical cues(*5*). Importantly, force sensing was proposed to contribute to infection, by inducing specific mechanical morphotypes with enhanced tolerance and invasiveness(*6*, *7*).

Despite these recent advances, how forces experienced by bacteria in their natural environment influence their physiology, morphology and growth remains poorly understood, especially in the context of disease progression during infection(*6–9*). Bacteria form dense multicellular communities in a wide range of conditions, for example during biofilm formation upon cell growth within a self-secreted polymeric matrix(*10*). The combination of bacterial cell proliferation, cell-cell cohesion and adhesion on a substrate was shown to induce the build-up of internal stress, suggesting that this might be a key general feature of biofilm growth(*9*). The geometry of the microenvironment can also impose external constraints on growing bacterial cells(*11*). More generally, bacterial growth in a limited space leads to the formation of dense confined colonies, leading to compressive forces. In eukaryotes, cells proliferating in a limited space generate pressures ranging from the kPa range for animal cells and up to the hundreds of kPa range in the case of yeast(*12–15*). These self-induced compressive forces affect the regulation of key cellular processes, from cell growth and division to differentiation and stress tolerance(*16*, *17*). However, the magnitude of growth-induced turgor pressure for bacterial colonies and its impact on bacterial physiology at the single-cell scale remain elusive(*11*). Historically, the bacterial cell wall was considered the sole element responsible for bearing the mechanical stress produced by turgor pressure and maintaining cellular shape. However, a growing appreciation of the mechanical role of the outer membrane in Gram-negative bacteria has emerged(*18*, *19*). Additionally, other envelope layers synthesized by bacteria in response to cell wall damage, such as the capsule, may be structurally important for mechanical cell integrity(*20*). Understanding which components are required in bacteria for mechanosensing and adaptation to confined growth might reveal novel mechanisms of adaptation to stress and specific molecular targets to fight bacterial infections.

Indeed, the potential impact of spatial confinement and the consequent generation of growth-induced mechanical pressure is particularly relevant for the infectious context, where bacteria grow within the host tissue and are potentially submitted to increasing mechanical constraints as the colony expands. For example, uropathogenic *Escherichia coli* (*E. coli*) invade cells of the bladder epithelium forming dense intracellular bacterial communities (IBCs)(*21*, *22*). These colonies can grow until the burst of the infected cell and the release of bacteria contributes to recurrent infections(*23*, *24*). Another example of pathogenic bacteria facing confinement during host invasion is *Neisseria meningitidis* growing and ultimately occluding the lumen of blood vessels(*8*). *Staphylococcus aureus* can also form very tight colonies of deformed bacterial cells in canaliculi of cortical bone. This leads to chronic osteomyelitis cases, a major challenge in orthopedics(*25*). Bacteria growing in a physically-limited environment is thus occurring in many infectious diseases, however further studies are required to dissect the functional impact of this process in disease progression.

Here, we combined *in vitro* and *ex vivo* experiments together with theoretical modeling to unveil how bacteria respond and adapt to spatial confinement. We developed a microfluidic system to precisely confine *E. coli* colonies while allowing live-cell microscopic observation at subcellular resolution. We found that spatial confinement induces profound changes in bacterial physiology. Growth under confinement increases intracellular crowding and turgor pressure. These modifications in cell physical properties influence growth and division in a way that impacts bacterial morphology and are driven by transcriptional changes in the bacterial transcriptional profile upon confinement. Specifically, the Rcs pathway leads to cell envelope remodeling essential to counterbalance high levels of turgor pressure and ensure bacterial shape maintenance in these conditions, both *in vitro* and during urinary tract infections in a mouse model. Overall, our work provides a physical and mechanistic elucidation of bacterial growth in confined space for commensal and pathogenic strains.

## RESULTS

### Bacterial proliferation under confinement generates growth-induced pressure

To investigate how bacteria adapt to mechanical confinement, we developed a PDMS-based microfluidic chip in which bacteria grow in space-limited chambers connected to 400 nm-wide nanochannels (**Figure 1.A**, **Supp. Figure 1.A**). The bacterial confiner was designed based on previous work performed on yeast and mammal cells and scaled down to dimensions adapted to trap 1 µm-wide cells(*12*, *14*). This setup allowed constant medium renewal while trapping bacteria inside the chambers (**Supp. Figure 1.B-C, Supp. Video 1**). Inside the bacterial confiner, bacteria proliferated, filled up the chambers in about 5 hours and became densely packed, *i.e.* confined (**Figure 1.B**). Bacterial death during extensive periods of confinement was rare (**Supp. Figure 1.D**). We asked whether bacterial proliferation upon confinement led to the generation of compressive forces by measuring PDMS chamber deformation. Semi-automatic tracking of the chamber contour showed that as soon as bacteria reached confluency, they pushed against the walls, leading to chamber deformation and physical confinement (**Figure 1.C**, **Supp. Figure 1.E-F, Supp. Video 2**). By contrast, in 5% of the cases, bacteria flowed out of the chambers at confluency without deforming the chamber. For quantification purposes, forces generated by bacteria were determined by calibrating PDMS deformability and single curves were aligned by defining time 0 as the time which precedes chamber deformation (**Supp. Figure 1.F-I**). We found that bacteria generated growth-induced pressure (GIP) averaging 300 kPa after about 10 hours of confined growth (**Figure 1.D**). 3D super-resolution imaging of *E. coli* K12 strain MG1655 with the inner membrane fluorescently labeled via a ZipA-mCherry fusion(*26*) showed that confinement did not induce any preferential 3D cell orientation, allowing quantification of bacterial features in 2D (**Supp. Figure 2.A-E**). Thus, 2D single-cell segmentation based on the inner membrane fluorescent marker was used to measure bacterial numbers in the 2D focal plane at the bottom of the chamber as a proxy for the number of bacteria in the chamber (**Supp. Figure 2.B**). We found that the number of bacteria continued to increase exponentially for 1.5 hours after confluency, which correlated with a progressive increase in growth-induced pressure (**Figure 1.D**, **Supp. Figure 1.I**). Thus, the bacterial confiner allowed for the first time dynamic subcellular observation of bacterial proliferation in a constrained space, while allowing efficient medium renewal and simultaneous pressure measurements(*11*). Using this device, we showed that bacterial proliferation upon confinement was sufficient to generate large compressive forces, an order of magnitude larger than the typical 1-10 kPa range of physiological forces found in mammalian eukaryotic tissues *in vivo*(*27*). This further raised the question of how bacteria generate these forces and adapt to mechanical confinement.

**Figure 1:**
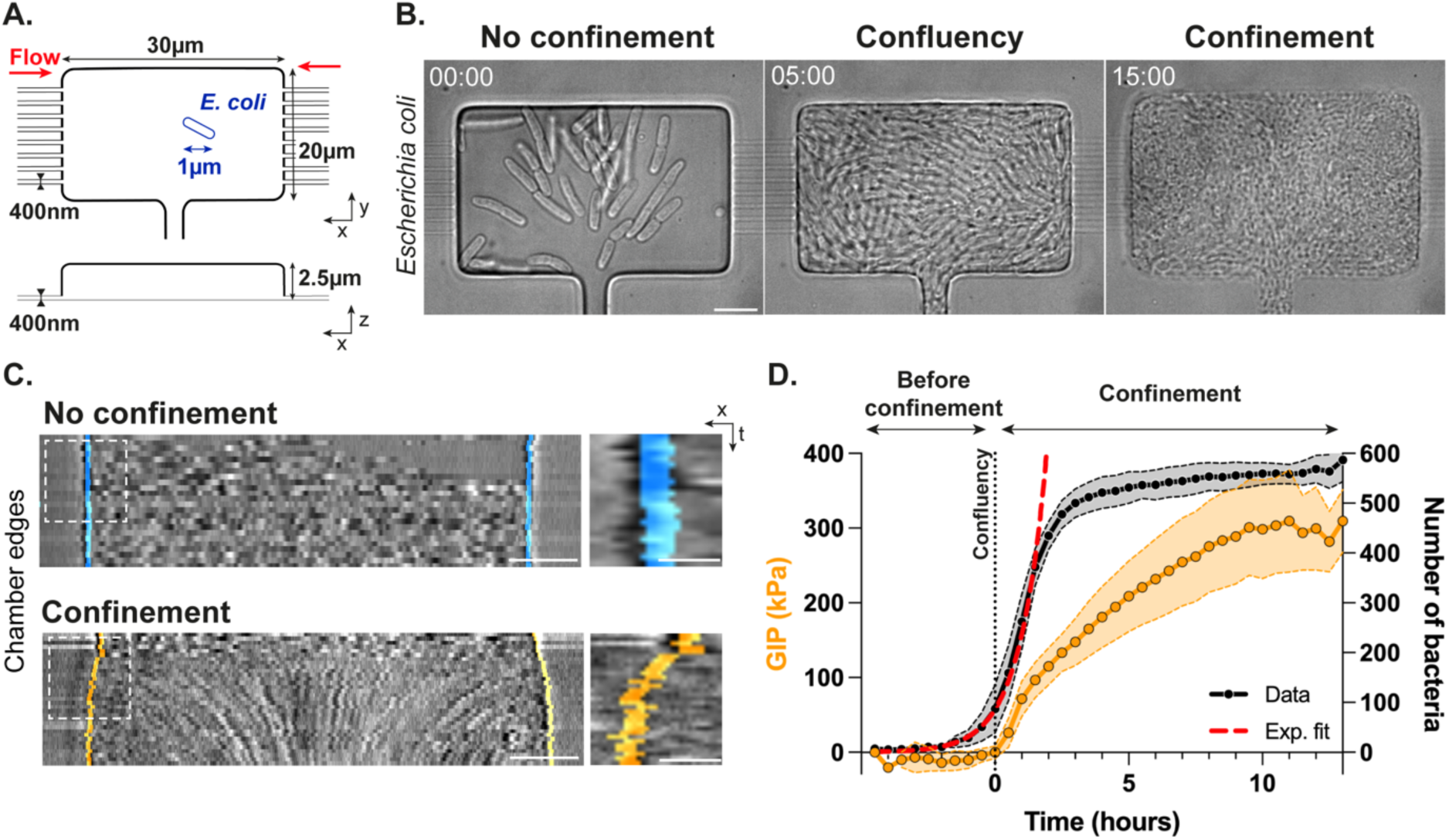
Bacterial proliferation under confinement generates growth-induced pressure. A. Schematics of the bacterial confiner in top *xy* and side *xz* views. Dimensions of the microfluidic chip are indicated respect to *E. coli* typical size (depicted in blue) inside the growth chamber. Medium renewal is ensured by flow passing in throughout the nanochannels as depicted by the red arrows. B. Timelapse brightfield images acquired at 30 minutes frame rate of bacterial proliferation in the bacterial confiner in the absence of confinement (t = 0 h, *left*), at confluency (t = 5 h, *middle*) and upon confinement (t = 15 h, *right*). Time is indicated as hh:mm. C. Kymographs of chamber edges for a control chamber where confluent bacteria proliferate without deforming the walls (*top*) and one chamber where bacterial proliferation upon confinement leads to chamber deformation along the x-axis (*bottom*). Segmented chamber edges are represented in color, blue and orange in the absence and presence of confinement respectively (see also **Supp. Video 2**). Inset: zoom in of the chamber deformation profile corresponding to the dashed white square regions. D. Growth-induced pressure (GIP, in kPa) and number of bacteria in the 2D focal plane of observation as a function of time (n_chambers_ = 13, N = 3). The phases before confinement, at confluency and upon confinement shown in Panel B are highlighted. Single curves are rescaled respect to the time 0 of pressure build-up. Data points correspond to mean values ± standard deviations. The number of bacteria follows an exponential growth curve until 1.5 hours (red dashed line). Scale bars: 5 µm, Scale bars insets: 1 µm.

### Growth-induced pressure leads to rod cell shortening due to uncoupling between bacterial growth and division

To decipher how mechanical confinement affects bacterial physiology, we characterized its impact on bacterial cell shape using a bacterial strain expressing the inner membrane marker described above (**Figure 2.A -*top*, Supp. Video 3**). Single cell area decreased upon confinement, as depicted in the colormap (**Figure 2.A - *bottom*, Supp. Video 3**). Average bacterial area rapidly dropped by a factor of 4 specifically at the onset of pressure build-up reaching a stable minimal area of 0.5 µm^2^ 2 hours later (**Figure 2.B, Supp. Figure 3.A-B**). By quantifying changes in bacterial length and width over time (**Supp. Figure 2.C**), we found that this morphological transition was mostly due to a 75% decrease in bacterial length (**Figure 2.C, Supp. Figure 3.C**). To determine whether this morphological transition was triggered by an uncoupling between growth and division, we imaged bacterial proliferation at higher temporal resolution to reconstruct single-cell lineages in the 2D plane of observation(*28*) (**Figure 2.D, Supp. Figure 2.D-E**). We found that while bacterial growth rate rapidly decreased at the onset of confinement, division rate persisted for a period of about 30 minutes. In other words, bacteria continued dividing for about 2 cell cycles while their growth was almost completely arrested, leading to cell shortening (**Figure 2.D, Supp. Figure 3.D**). The division rate then decreased during the following 90 minutes, reaching a state where bacteria stopped growing and dividing. These data show that in *E. coli*, growth-induced pressure initially caused a loss of size control through the persistence of division, as previously reported for mammalian cells in a confluent monolayer(*29*). Since fresh medium was continuously provided, these changes were not due to starvation, and the size reduction we observed was stronger than starvation-induced shortening of bacteria (**Figure 2.E**). Our results identified 3 phases that followed confinement, defined as the onset of growth-induced pressure: (1) bacterial growth declined rapidly while division persisted, leading to a decrease in length (**Figure 2.E**); (2) division rate decreased and finally (3) both growth and division rates were arrested while growth-induced pressure kept increasing (**Figure 2.D**).

**Figure 2:**
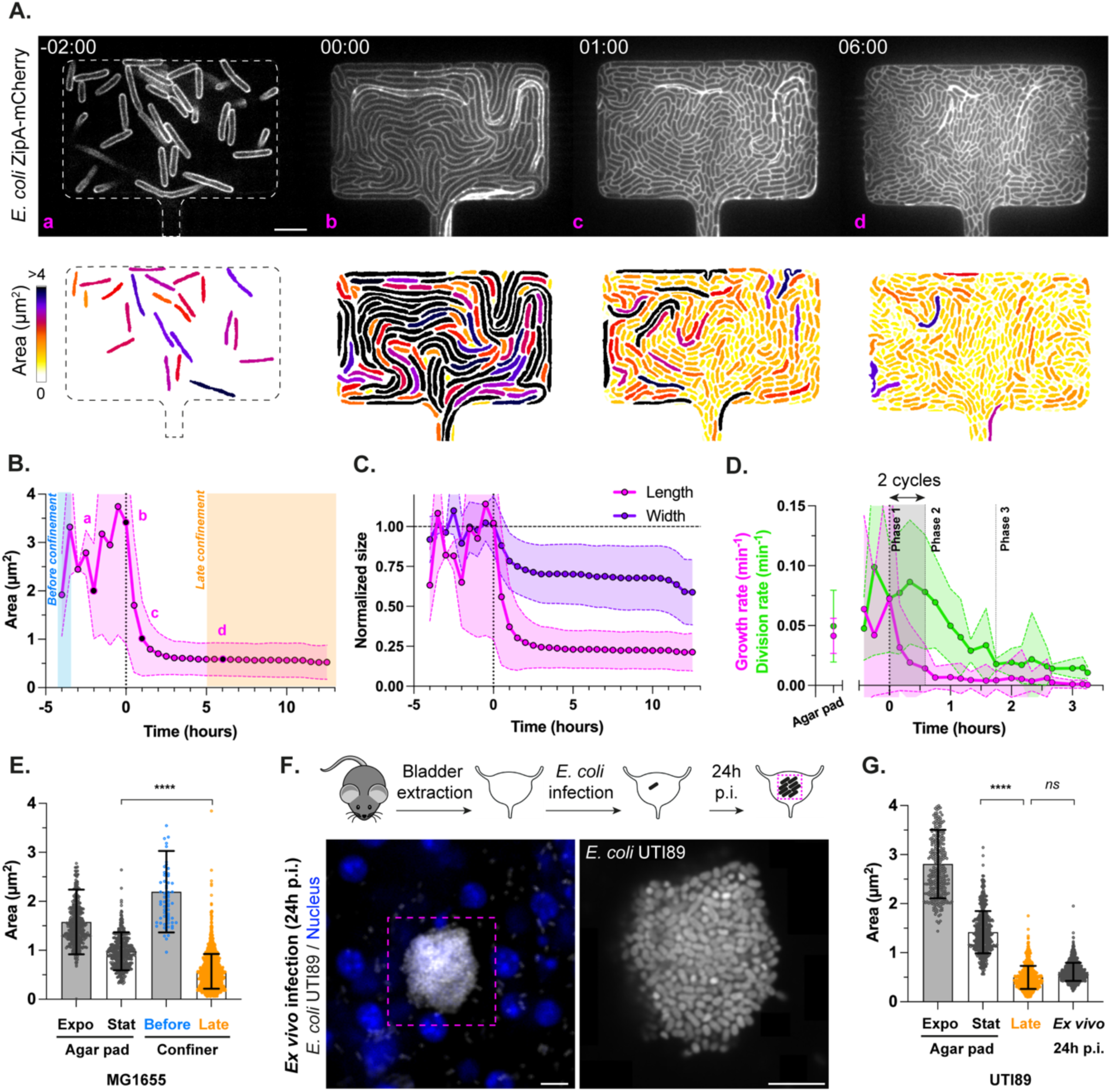
Growth-induced pressure leads to rod cell shortening due to uncoupling between bacterial growth and division. A. Timelapse confocal images acquired at 30 minutes frame rate of *E. coli* MG1655 ZipA-mCherry bacteria fluorescently labeled at the inner membrane during proliferation in the bacterial confiner upon confinement (*top*). Corresponding colormap of the single-cell bacterial area (*bottom*). The phases before confinement (t = - 2 h), at the time at which pressure builds up (t = 0 h), 1 h and 6 h after pressure build-up are shown in a, b, c and d respectively. Time is indicated as hh:mm (see also **Supp. Video 3**). B. Quantification of individual bacterial area over time (n_bacteria_ > 47000, n_chambers_ = 4, N = 2). Data points correspond to mean values ± standard deviations. Letters refer to the phases depicted in Panel A. The regions highlighted in colors refer to the datasets used for the categories Before and Late confinement in Panel F. C. Quantification of bacterial length and width normalized by their average value before confinement (n_bacteria_ > 47000, n_chambers_ = 4, N = 2). Data points correspond to mean values ± standard deviations. D. Quantification of single-cell growth and division rate over time, obtained from single-cell tracks at 5 minutes frame rate. The time interval corresponding to Phase 1 during which the uncoupling takes place because of two complete division cycles in the absence of growth is highlighted in grey (n_bacteria_ = 26728, n_tracks_ > 800, n_chambers_ = 2, N = 1). E. Quantification of individual bacterial area for MG1655 strain in different conditions: exponential (n_bacteria_ = 482, N = 3) and stationary phases (n_bacteria_ = 410, N = 3) in agar pad, before (n_bacteria_ = 65, n_chambers_ = 4, N = 2) and upon late confinement (n_bacteria_ = 1721, n_chambers_ = 4, N = 2) in the bacterial confiner. F. Schematics of the main steps of *ex vivo* mouse bladder infection by uropathogenic *E. coli* (*top*). Corresponding confocal images of an infected uroepithelium at 24 h post-infection (p.i.) showing tight aggregates of the uropathogenic *E. coli* UTI89 strain confined within the tissue (grey: bacteria; blue: nuclei) at low (*bottom left*) and high resolution (*bottom right*). G. Quantification of individual bacterial area for UTI89 strain in different conditions: exponential (n_bacteria_ = 318, N = 3) and stationary phases (n_bacteria_ = 542, N = 3) in agar pad, upon late confinement (n_bacteria_ = 701, n_chamber_ = 1, N = 1) in the bacterial confiner and 24 h post-infection (n_bacteria_ = 1096, n_aggregates_ = 11, N = 4) are indicated. Data points correspond to mean values ± standard deviations. Statistical significance of the results was assessed using Welch ANOVA tests. All scale bars: 5 µm.

We then wondered whether these morphological changes observed in PDMS chambers also occurred during infection (**Figure 2.F**). We quantified individual bacterial areas of a patient-derived uropathogenic *E. coli* (i.e., UPEC) strain, UTI89(*30*), under agar pads, in the bacterial confiner, and in intracellular bacterial communities (i.e., IBCs) formed during *ex vivo* infection of mouse uroepithelium. UPEC showed size reductions similar to the K12 strain, both in the confiner and inside urothelial cells, highlighting the clinical relevance of our observations (**Figure 2.F-G, Supp. Figure 3.E-F**). These morphological changes were reversible upon release of the pressure in the chambers (**Supp. Figure 3.G-H, Supp. Video 4**), which is reminiscent of the morphological plasticity described in urinary tract infections (i.e., UTIs) upon cell rupture and bacterial release(*23*).

### Confinement increases cytoplasmic crowding through changes in protein concentrations and DNA occupancy

We next explored the impact of confinement on bacterial physiology at the subcellular scale. We hypothesized that growth-induced pressure may impact the production and accumulation of two major components of the bacterial cytoplasm: proteins and DNA. At the protein level, we investigated the impact of pressure build-up using two fluorescent reporters, one under the control of a constitutive promoter (P*_R_*-meGFP) to have a global readout of protein transcription and another with the promoter of the *ftsZ* gene, a key component of the division machinery (P*_ftsZ_*-FtsZ-mNeonGreen(*31*)) (**Figure 3.A, Supp. Figure 4.A**). We found that in both cases, cytoplasmic mean fluorescence intensities, proportional to protein concentration, increased in the two first phases of confinement (**Figure 3.B - *top*, Supp. Figure 4.B-C, Supp. Video 5**). By computing the protein production rate, we found that this was because synthesis of both GFP and FtsZ declined at the onset of confinement more slowly than the cell growth rate (**Figure 3.B - *bottom*, Supp. Figure 4.D-E**). This shows a non-specific increase in protein concentration in the cytoplasm of confined bacteria during Phases 1 and 2. At the DNA level, we explored whether the initial persistence of bacterial division in the absence of growth could lead to a higher DNA occupancy in the cytoplasm. For this, we used a HU-GFP fusion to measure the karyoplasmic or N:C ratio, defined as the ratio between DNA and cell area(*32*) (**Supp. Figure 2.C**). We found that confined bacteria exhibited a 20% increase in karyoplasmic ratio during the two first phases of confinement compared to control, non-confined bacteria (**Figure 3.C-D, Supp. Figure 4.F-G, Supp. Video 5**). This meant that pressure build-up induced the formation of small bacteria with a cytoplasm largely occupied by the nucleoid suggesting that growth-induced pressure induces higher levels of cytoplasmic crowding in bacteria, as previously shown in eukaryotes(*13*, *14*). To evaluate cytoplasmic apparent viscosity, we expressed 40nm-wide diffusive genetically encoded nanoparticles (40 nm-GEMs) and tracked them in the bacterial cytoplasm(*33*) (**Figure 3.E, Supp. Figure 4.H, Supp. Video 5**). In line with our previous results, we found that 40nm-GEMs diffusion slightly decreased in the first phase of confinement and more steeply in the second phase (**Figure 3.F**), showing that cytoplasmic crowding significantly increased upon confinement. Collectively, these results demonstrated that growth-induced pressure was associated with a highly crowded bacterial cytoplasm mediated by protein and DNA accumulation.

**Figure 3:**
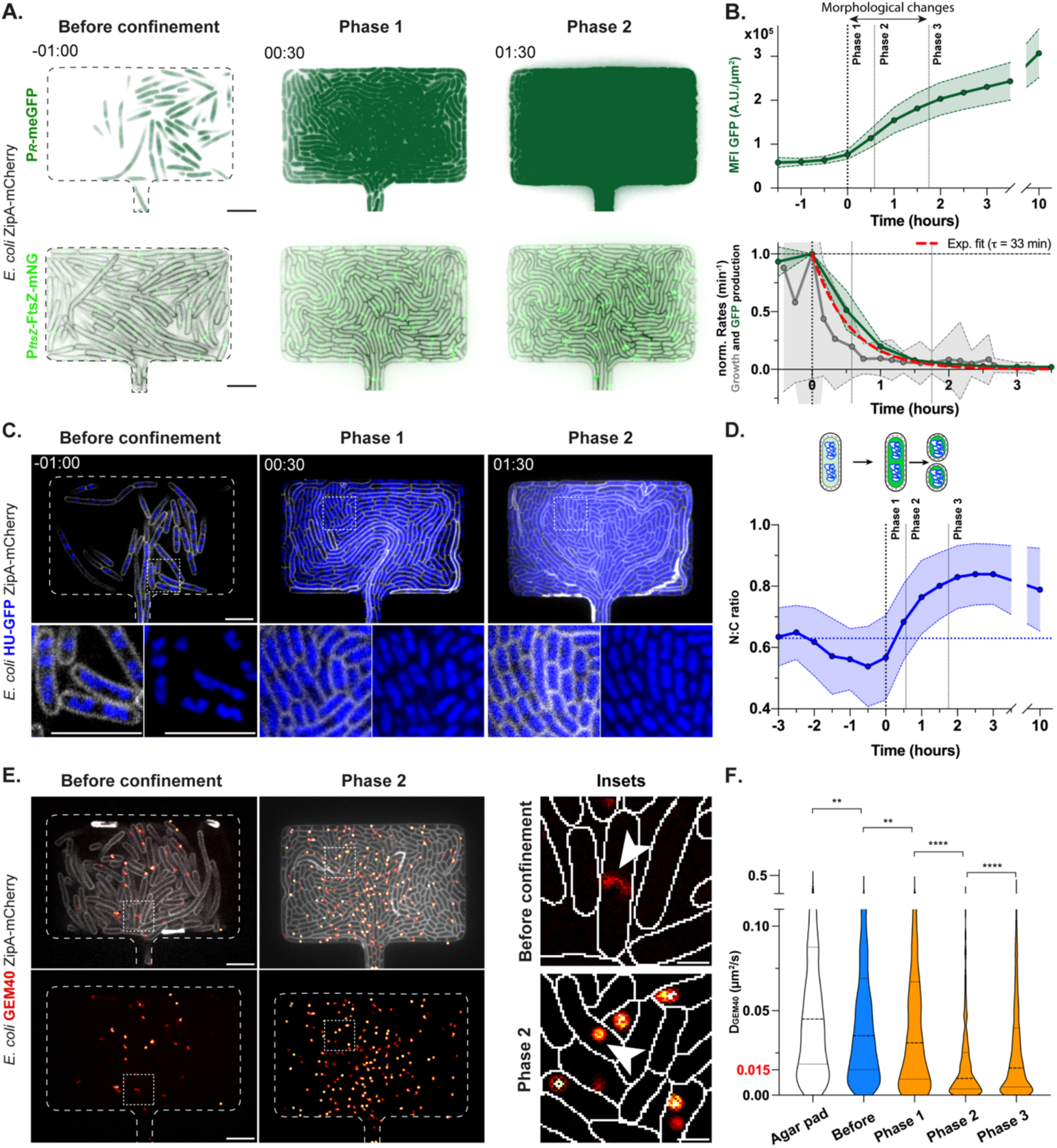
Confinement increases cytoplasmic crowding through changes in protein concentrations and DNA occupancy. A. Timelapse confocal images acquired at 30 minutes frame rate of *E. coli* MG1655 GFP ZipA-mCherry bacteria expressing in their cytoplasm a constitutive GFP and proliferating in the bacterial confiner in the presence of confinement (*top*). Similar timelapse of *E. coli* BW27783 FtsZ-mNG ZipA-mCherry bacteria expressing the fusion protein FtsZ-mNG (*bottom*). The two signals are indicated in dark and light green respectively. Bacterial cell contour is detected with the ZipA-mCherry signal as in Figure 2, and hereby indicated in black when necessary. Representative examples of the phases before confinement, 1 and 2 are shown. Time is indicated as hh:mm (see also **Supp. Video 5**). B. Temporal evolution of single-cell mean fluorescent intensity of the constitutive GFP (n_bacteria_ = 50661, n_chambers_ = 4, N = 1) signal over the entire cell area respect to pressure build-up, indicated in dark green (*top*). Phases 1, 2, 3 induced upon confinement are highlighted, together with the time period characterized by major morphological changes. Quantification of total GFP (n_chambers_ = 4, N = 1) production rate normalized by its value at time 0 of pressure build-up, and corresponding exponential fit (typical time τ = 33min), indicated in dark green and dashed red respectively (*bottom*). Bacterial growth rate is indicated in grey and computed on another dataset, as shown in Figure 2.D. Data points correspond to mean values ± standard deviations. C. Timelapse confocal images acquired at 30 minutes frame rate of *E. coli* HU-GFP ZipA-mCherry bacteria fluorescently labeled at the inner membrane (grey) and DNA (blue) proliferating in the bacterial confiner upon confinement (*top*). Insets zoom in the white dashed square regions and highlight DNA occupancy in the bacterial cytoplasm (*bottom*). Representative examples of the phases before confinement, 1 and 2 are shown. Time is indicated as hh:mm (see also **Supp. Video 5**). D. Schematics of the hypothesis of increase in protein concentration (indicated with a shift from light to dark green) and DNA occupancy (blue) upon confinement and rod shortening (*top*). Quantification of the karyoplasmic ratio (indicated as the ratio between the bacterial cell and nucleoid areas) over time regarding pressure build-up (n_bacteria_ = 26894, n_chambers_ = 2, N = 1) (*bottom*). E. Confocal images of *E. coli* GEM40 ZipA-mCherry bacteria expressing GEM40 diffusive nanoparticles in their cytoplasm while proliferating in the bacterial confiner in the absence of confinement (*left*) or in Phase 2 of confinement (*middle*). Bacterial cell contour is detected with the ZipA-mCherry signal as in Figure 2, hereby indicated in grey. Streaming acquisition at 50 ms frame rate of GEM40 diffusion allowed to generate single particle tracks, here indicated in color. The colormap represents the duration of the nanoparticle tracks, red corresponding to short trajectories and white to longer ones. Insets correspond to the white dashed square regions and zoom in individual GEM40 trajectories within the bacterial cytoplasm drawn based on the membrane signal (*left*) in the absence of confinement (*top*) or in Phase 2 of confinement (*bottom*). White arrows indicate one typical GEM40 trajectory for each condition. Representative examples of the phases before confinement and 2 are shown. Scale bars insets: 1 µm (see also **Supp. Video 5**). F. Quantification of the GEM40 effective diffusion coefficient in different conditions: in agar pad (n_tracks_ = 1020, N = 2), before confinement (n_tracks_ = 1120, N = 3) and upon confinement in Phase 1 (n_tracks_ = 444, N = 3), Phase 2 (n_tracks_ = 1431, N = 3), and Phase 3 (n_tracks_ = 2517, N = 3). The diffusion value of 0.015 µm^2^/s corresponding to the higher level of crowding reached upon confinement is highlighted in red. Datasets are represented as violin plots with highlighted median values and corresponding quartile range. Statistical significance of the results was assessed using Kruskal-Wallis tests. All scale bars except the insets in Panel E: 5 µm.

### A progressive increase in cytoplasmic crowding is sufficient to recapitulate the bacterial division trend observed upon confinement

We next asked by which mechanisms bacteria modulate division upon confinement, in particular the role of the observed increase in cytoplasmic crowding. We hypothesized that a progressive increase in crowding could first lead to division induction and then division arrest. Previous studies suggest that the accumulation of proteins of the divisome is sufficient to trigger bacterial division, both in steady-state conditions and upon hyperosmotic shock(*34*, *35*). By tracking 40nm-GEMs particles in bacterial cells subjected to a range of sorbitol concentrations, we confirmed that hyperosmotic shock led to an increase in cytoplasmic crowding (**Figure 4.A**). To mimic a progressive increase in crowding, we then performed two successive hyperosmotic shocks at increasing sorbitol concentrations (0.5M and 1M) and monitored bacterial divisions. We observed that while bacteria often divide a few minutes after the first shock, they rarely divide after the second shock (**Figure 4.B, Supp. Figure 5.A-B**). A progressive increase in osmotically-induced crowding thus reproduced the confinement-induced change in bacterial division characterized by a first increase in the fraction of dividing cells followed by a sharp decrease (**Figure 4.C-D**). These results suggested that, while an initial increase in protein concentration was sufficient to trigger bacterial division, an additional increase in intracellular crowding rather inhibited bacterial division. Interestingly, despite high karyoplasmic ratios in confined bacteria (**Figure 3.C-D**), the reduction in division rate was not regulated by the nucleoid occlusion regulator SlmA(*36*) (**Supp. Figure 5.C-E**). Therefore, these data support that cytoplasmic crowding regulates bacterial division upon confinement through physical means, by preventing protein diffusion and/or protein synthesis as previously proposed in yeast(*13*), ultimately leading to the formation of non-growing and non-dividing highly crowded bacteria. In this state, confined bacteria seem to have reached a quasi-frozen stalled state, in terms of division and protein synthesis. Nevertheless, unexpectedly, we observed that the growth-induced pressure kept increasing steadily for hours after both protein synthesis and division had stopped. We proposed to name this defined third phase of confined growth *overpressurization* regime.

**Figure 4:**
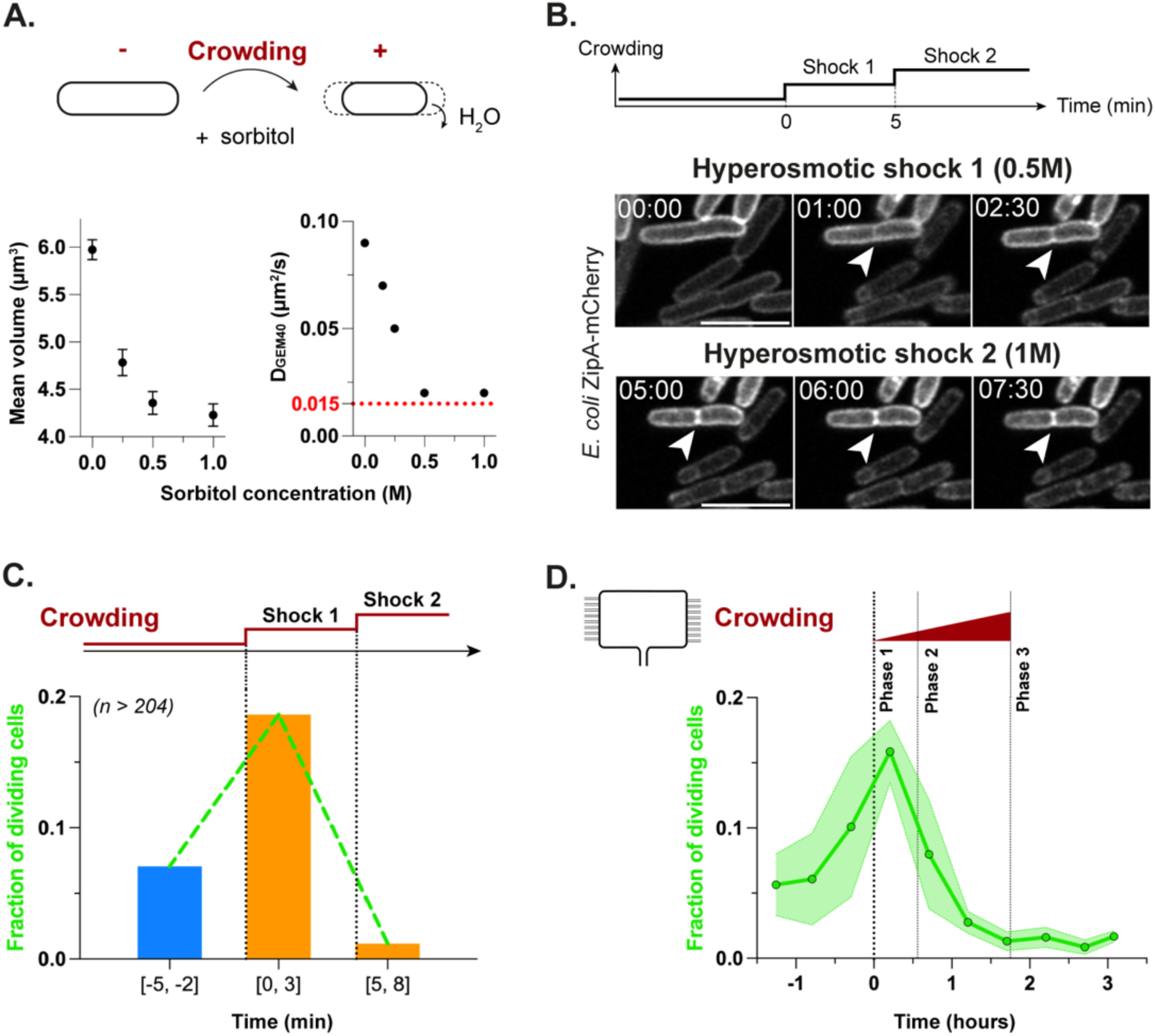
A progressive increase in cytoplasmic crowding is sufficient to recapitulate the bacterial division trend observed upon confinement. A. Schematics of one *E. coli* bacterium submitted to a hyperosmotic shock by increasing the osmolarity of the culture medium using sorbitol in a classical flow assay (*top*). Water flowing out of the cell leads to a decrease in bacterial size and a subsequent increase in crowding, highlighted in dark red. Quantification of the mean bacterial volume (*bottom left*) and GEM40 effective diffusion coefficient (*bottom right*) for a range of sorbitol concentrations (for each condition: n_bacteria_ > 130, N > 3). The diffusion value of 0.015 µm^2^/s corresponding to the higher level of crowding reached upon confinement is highlighted in red. Data points correspond to mean values ± standard deviations. B. Schematics of the increase in crowding induced upon two successive hyperosmotic shocks, a first one of 0.5 M sorbitol at time 0, and a second one of 1 M sorbitol at time 5 minutes (*top*). Timelapse confocal images acquired at 1 minute frame rate of *E. coli* ZipA-mCherry fluorescently labeled at the inner membrane submitted to the two successive hyperosmotic shocks (*bottom*). White arrows point at the bacterial division site. Time is indicated as mm:ss. Scale bars: 5 µm. C. Quantification of the fraction of dividing cells in response to two successive hyperosmotic shocks (per condition: n_bacteria_ > 204, N = 4). D. Quantification of the fraction of dividing cells upon confinement, with a highlight on Phases 1, 2 and 3 (n_bacteria_ = 26728, n_tracks_ > 800, n_chambers_ = 2, N = 1). To note, the increase in the fraction of dividing cells slightly precedes the onset of GIP build-up likely because of a lack of accuracy in the estimation of this critical timepoint, the deformation of the chamber being acquired at a lower temporal resolution than bacterial division.

### Theoretical modeling reveals a central role for cell anabolism in the *overpressurization* regime

Although the mechanistic origin of turgor pressure might differ between organisms, it is often assumed that an arrest in protein synthesis should also stop growth-induced pressure(*13*, *37*). To understand our non-trivial observation of steady pressure increase in confined bacterial colonies, we built a minimal theoretical model of bacterial growth that leverages recent knowledge in mechano-osmotic regulation(*38–40*) (**Supp. Model**). We modelled confining chambers with an initial rectangular shape that deformed elastically in 3D due to the mechanical pressure exerted by bacterial growth. Average bacterial cell shape was approximated by cylinders (**Figure 5.B**). Bacterial cell volume is determined by the balance between the osmotic pressure and the mechanical pressure difference throughout the bacterial envelope(*41*). Before confinement onset, an increase in the intracellular osmotic pressure, due to growth-mediated accumulation of trapped osmolytes, results in an increase in cell volume. Once bacteria fill up the entire space in the chamber, the intracellular osmotic pressure starts to be balanced by the chamber walls. In this regime, assuming homogenous bacterial density and behavior (**Supp. Figure 2.A**), deformation of the PDMS chamber walls (the growth-induced pressure) provides a direct measurement of the quantity of trapped osmolytes inside bacteria while remaining small compared to the total volume of the chamber (**Supp. Model**). Because proteins initially keep being produced at a normal rate while cell volume increase is constrained by the chamber, protein concentration increases. We incorporated in the model the notion that bacterial division is triggered at a threshold number of divisome proteins(*34*), imposing that *in silico* bacteria divide once their protein number has been doubled, leading to smaller cells as observed experimentally. This describes the first phase of confinement, during which the bacterial growth in volume rapidly stalls, constrained by the walls of the chamber, while several rounds of division produce smaller cells. To account for the second phase, we incorporated the now well-established notion that, due to the size of abundant protein complexes such as ribosomes, protein accumulation is accompanied by cytoplasmic crowding, leading to a decrease in protein production rate, as observed experimentally(*13*). This is described in the model by a simple equation that, instead of assuming a constant protein accumulation rate, couples it to the global protein concentration, similarly to several recent studies in yeast and other organisms(*13*, *37*). Prior models would assume that, at the end of the second phase, once cytoplasmic crowding has led to an arrest of protein accumulation, growth-induced pressure should also stop increasing. Because we instead observed a persistent pressure increase, we introduced a key novel feature in the theoretical framework, based on recent work aimed at explaining how growth in volume (accumulation of intracellular trapped osmolytes) and protein accumulation can be decoupled, leading to cytoplasmic dilution in overgrowing yeast cells(*38*, *42*). We indeed reasoned that, similarly to this prior work, there is a decoupling of two types of osmolytes in the third phase post-confinement (**Figure 5.B**): (1) small osmolytes, e.g. metabolites and counterions, which were recently proposed to dominate the intracellular trapped osmolytes, due to their large numbers inside cells (*43*); (2) proteins, whose contribution is mostly steric and whose production rate is reported to be sensitive to cytoplasmic crowding(*13*), but, due to their low numbers, have a minor direct contribution to intracellular osmotic pressure.

**Figure 5:**
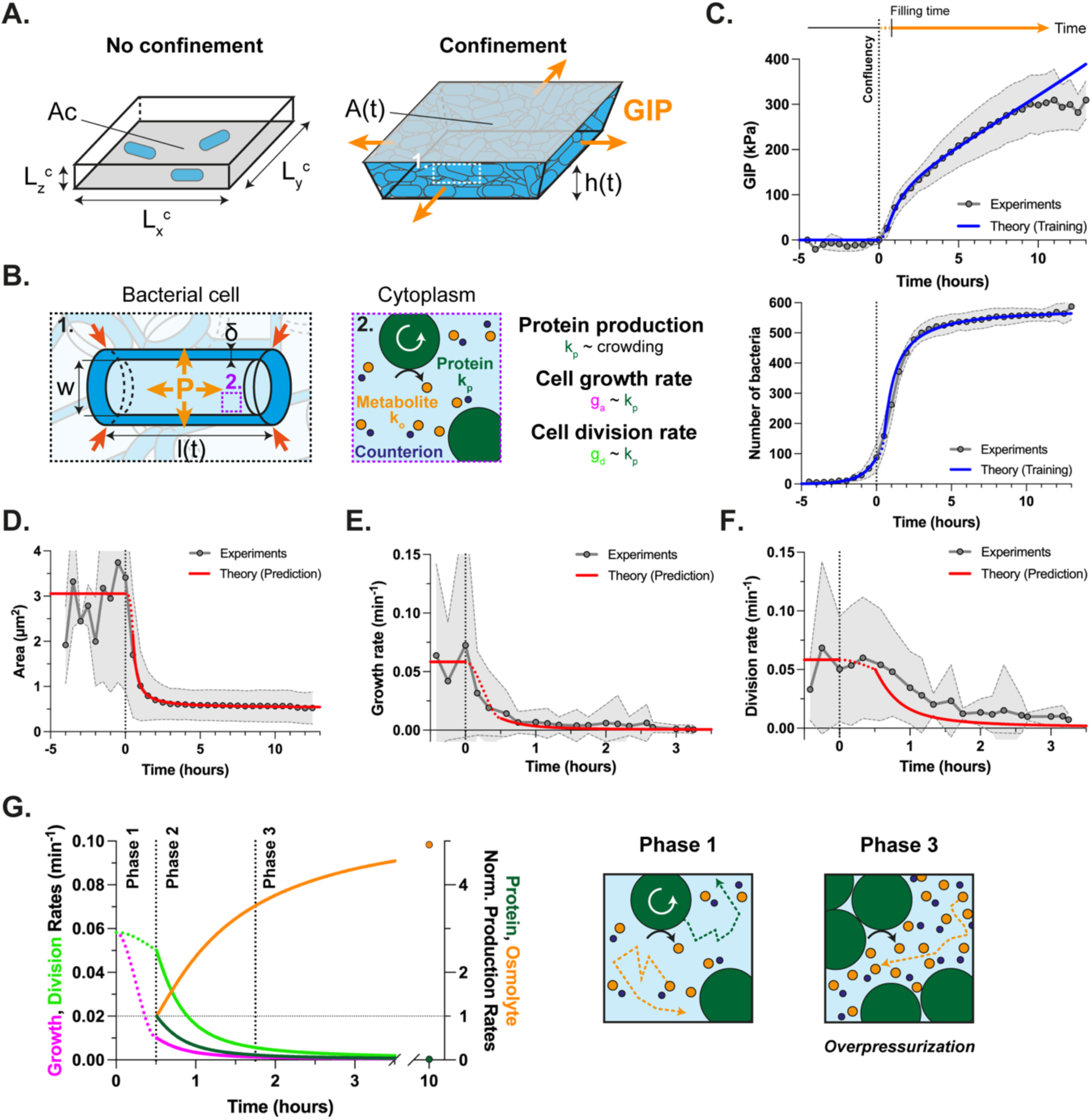
Theoretical modeling reveals a central role for cell anabolism in the *overpressurization* regime. A. Schematics of the theoretical model for bacterial growth in a limited space. The microfluidic chamber is modeled as a rectangular 3D space of dimensions L_x_^c^ = 20 µm, L_y_^c^ = 30 µm, L_z_^c^ = 2.6 µm and projected area A_c_ (*left*), which deforms elastically due to the mechanical pressure exerted by bacterial growth (*right*). B. Bacteria are modeled as cylinders of width w and length l(t) enclosed in a shell δ corresponding to the cell envelope (*left*). The osmotic pressure is defined by the production of osmolytes, including small osmolytes (e.g. metabolites and counterions, whose synthesis depends on proteins as depicted by the black curved arrow) and proteins (*right*). Bacteria are characterized by a crowding-dependent protein production rate (denoted k_p_), that in turn defines bacteria growth and division rates (denoted g_a_ and g_d_ respectively). C. Bacterial growth upon confinement is modeled starting at time t_0_, defined as the time where 2D confluency is reached (*top*). To note, the theoretical value of t_0_ (called Filling time) is delayed of 30 minutes compared to the experimental one (written Confluency) (**Supp. Figure 6.A-B**). For this reason, during the first 30 minutes, the model interpolates the bacterial features and is represented with dotted lines. Experimental curves (grey) and theoretical fits (blue) of growth-induced pressure (*top*) and the number of bacteria in 2D (*bottom*) as a function of time, both curves being used to train the model and fix the adjustable parameters. D. Theoretical prediction (red) and corresponding experimental curve (grey) of the temporal evolution of bacterial area upon confinement. E. Theoretical prediction (red) and corresponding experimental curve (grey) of the temporal evolution of bacterial growth rate. F. Theoretical prediction (red) and corresponding experimental curve (grey) of the temporal evolution of bacterial division rate (expressed as ln(2)*experimental division rate, see **Supp. Model**). G. Theoretical predictions of bacterial growth rate (magenta), division rate (light green), protein normalized production rate (dark green), and osmolyte normalized production rate (orange) (*left*). The model allows to identify and characterize the regimes of bacterial confinement corresponding to Phase 1, 2 and 3. A schematics of Phase 1 and 3 illustrating the differential impact of crowding on proteins and small osmolytes is represented (*right*). In Phase 1, both proteins and small osmolytes are produced (as indicated by the white and black arrows respectively) and freely diffuse in the cytoplasm (as indicated by the dotted green and orange arrows respectively). Note that in Phase 3, bacteria are non-growing and non-dividing due to crowding but still produce osmolytes, leading to *overpressurization* of the bacterial cytoplasm.

This model accounts for the three phases of confined growth based on a simple force balance complemented by three ingredients: 1) cell division depends on the doubling of protein number; 2) protein accumulation depends on protein concentration with a saturation effect due to crowding, as described before(*13*); 3) intracellular trapped osmolytes, responsible for the growth-induced pressure, are dominated by small biomolecules produced by proteins, with an accumulation rate that depends on the anabolic activity of cells that we simply assume to be proportional to the total number of proteins in the cell.

Because our model relies on a small number of explicit parameters with observables that can be directly measured experimentally, we were able to test it quantitatively. We first fitted three parameters using experimental growth-induced pressure measurements and the number of bacteria at the bottom of the chamber (**Figure 5.C**, **Supp. Model**). We then produced predictions without any adjustable parameter on the other independent datasets at our disposal: namely bacterial area, growth and division rate, and diffusion of 40 nm-GEMs particles (**Figure 5.D-F**, **Supp. Figure 6.E**). The quantitative agreement between theoretical predictions and experimental measurements validated our model. This allows us to define the different regimes of bacterial proliferation under confinement in physical terms.

A key feature of the model is that growth-induced pressure increases for hours, even in the absence of protein production, as observed experimentally (**Figure 5.C**, **Supp. Figure 6.F**). This arose from a fundamental difference between the types of osmolytes modeled here: proteins promote their own production while small osmolytes, mostly metabolites and counterions, rely on proteins for production, import, or degradation. Therefore, in the limiting regime of constant protein number, the model predicts a linear increase in the number of trapped osmolytes, resulting in a linear increase of the osmotic pressure and thus of the growth-induced pressure, as observed experimentally in the first hours of the third confinement phase (**Figure 5.G**). Of note, we verified that the predictions of the model are not altered if we consider that anabolic activity is also reduced due to protein crowding (**Supp. Model**, **Supp. Figure 6.G-J**). The late saturation in growth-induced pressure could have a variety of causes, such as passive or active shut down of anabolic activity. It is not directly explained by our model and could correspond to a fourth phase of confinement, which is out of the scope of this study. Thus, by decoupling growth and essential biological processes such as proteins and small osmolytes production, mechanical confinement induced a unique bacterial state during which bacteria underwent an increase in their osmotic pressure, a situation that can be referred to as *overpressurization*. In this regime, our model correctly predicts the production of very elevated pressures within a few hours, reaching hundreds of kPa, despite a global arrest in protein synthesis and bacterial division. An important contribution of the model, beyond providing a convincing mechanistic explanation for our observations, is to suggest the importance of cell metabolism to produce a persistent pressure increase in confined bacteria, reaching levels that could potentially deform mammalian tissues in an infectious context.

### Rcs transcriptional response to mechanical confinement is required for shape maintenance in the *overpressurization* regime

Because uropathogenic *E. coli* experience confined growth during infection, we reasoned that targeting mechanosensory pathways required for the adaptation of bacteria to confinement might provide novel targets to reduce their virulence. By analogy to well-known mechanosensors in eukaryotes, we hypothesized that adaptation to elevated turgor pressure may rely on the bacterial envelope. We focused on the Regulator of Capsule Synthesis (Rcs) and Cpx pathways as they sense defects at the outer and inner membrane/peptidoglycan respectively(*44*, *45*). In addition, the mechanosensory ability of Rcs has recently been proposed(*20*, *46*). To assess the activation of the Rcs pathway upon confinement, we used the P*_rcsA_*-GFP transcriptional reporter(*47*) (**Supp. Figure 7.A**). We found that the Rcs pathway was activated at the onset of confinement as soon as the pressure built up in the chambers (**Figure 6.A-B**, **Supp. Figure 7.B, Supp. Video 6**). Similarly, the Cpx envelope stress response was transcriptionally activated upon confinement, whereas the RecA pathway, induced upon DNA damage, was not (**Supp. Figure 7.A, C-E, Supp. Video 6**). To decipher the role of the Rcs pathway in bacterial adaptation to confinement, we used a bacterial strain deficient in the Rcs response regulator *rcsB*. Strikingly, we found that, while the *rcsB* mutant exhibited a normal rod shape in the absence of confinement (**Supp.** Figure 7**.F**), it lost its shape at the onset of the *overpressurization* regime and progressively inflated at the edges of the chambers at the later stages of confinement (**Figure 6.C**, **Supp.** Figure 7**.G, Supp. Video 7**), reaching unusually large cellular surface areas (**Figure 6.D**). The *rcsB* mutant morphological changes were most striking in the areas where transcription of the Rcs regulon was the highest (**Figure 6.B**). By contrast, a *cpxR* mutant deficient in the Cpx envelope stress response displayed only classical rod shape morphologies (**Supp. Figure 7.H, Supp. Video 7**), showing that the Rcs pathway specifically played a key role in shape maintenance upon late confinement.

**Figure 6:**
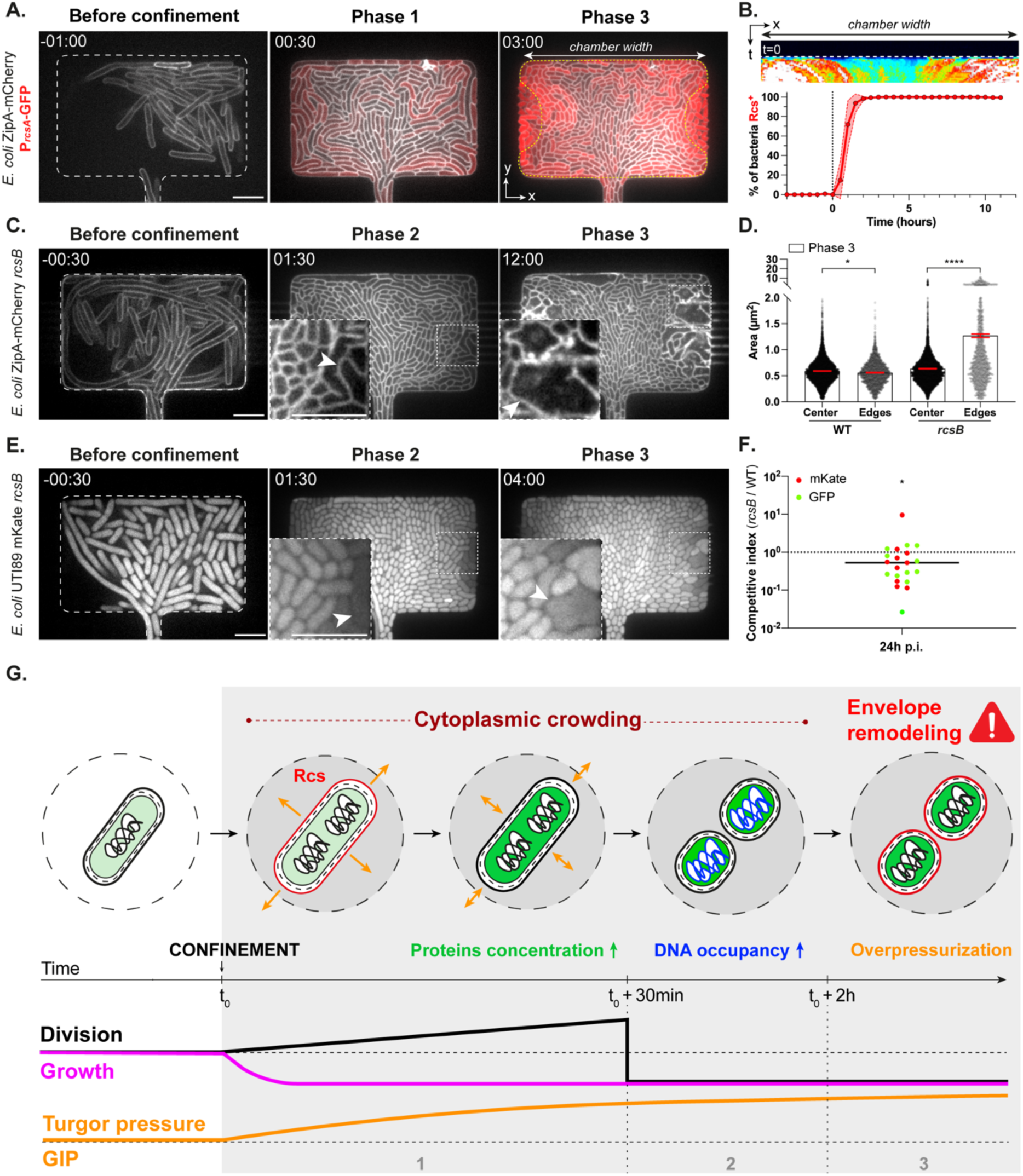
Rcs transcriptional response to mechanical confinement is required for shape maintenance in the *overpressurization* regime. A. Timelapse confocal images acquired at 30 minutes frame rate of a *E. coli* MG1655 ZipA-mCherry P*_rcsA_*-GFP strain fluorescently labeled at the inner membrane (grey) and expressing the Rcs transcriptional reporter P*_rcsA_*-GFP (red) during proliferation in the bacterial confiner upon confinement. Representative examples of the phases before confinement (-1 h), 1 and 3 (30 min and 3 h after pressure build-up respectively) are shown. The spatial pattern of the reporter fluorescence intensity is illustrated by a yellow dotted line (see also **Supp. Video 6**). B. Kymograph along the x-axis of P*_rcsA_*-GFP induction upon confinement, indicated with a Fire color scale (*top*). Quantification of the percentage of bacteria which activate the Rcs stress response (*bottom*). Time 0 indicates the time at which pressure builds-up in the chamber. Data points correspond to mean values ± standard deviations (n_bacteria_ = 35200, n_chambers_ = 3, N = 1). C. Timelapse confocal images acquired at 30 minutes frame rate of a *E. coli* ZipA-mCherry *rcsB* mutant deficient in the Rcs response fluorescently labeled at the inner membrane during proliferation in the bacterial confiner. Representative examples of the phases before confinement (-0.5 h), 2 and 3 (1.5 h and 12 h after pressure build-up respectively) are shown. Insets zoom in the regions depicted with a white dashed square line. White arrows indicate bacteria with morphological defects (see also **Supp. Video 7**). D. Quantification of bacterial area in Phase 3 upon confinement for the wild-type strain and the *rcsB* mutant either at the center or at the edges or the chamber. The regions named “Center” and “Edges” refer to the spatial pattern determined using the P*_rcsA_*-GFP fluorescent profile depicted in Panel A. Means ± standard errors are represented (per condition: n_bacteria_ ≥ 3928, n_chambers_ ≥ 2, N = 1). Statistical significance of the results was assessed using one-way ANOVA tests. E. Timelapse confocal images acquired at 30 minutes frame rate of a *E. coli* UTI89 mKate *rcsB* mutant deficient in the Rcs response fluorescently labeled in the cytoplasm during proliferation in the bacterial confiner upon confinement. Representative examples of the phases before confinement (-0.5 h), 2 and 3 (1.5 h and 4 h after pressure build-up respectively) are shown. Insets zoom in the regions depicted with a white dashed square line. White arrows indicate bacteria with morphological defects. F. *In vivo* competition experiment in a mouse model of urinary tract infection. Infections were performed by mixing two pairs of UTI89 bacterial strains, WT and *rcsB*, each of those in the same background, expressing GFP and mKate as indicated in green and red respectively. Competitive index 24 h p.i. (represented as median value) was calculated by dividing CFU/bladder of *rcsB* versus WT strains after tissue dissection and plating on corresponding selective antibiotics. Importantly, decreased fitness was observed using both pairs of strains, providing strong evidence for a role of *rcsB* in infection. Data obtained from all competition experiments were pooled together for calculation of the competitive index and statistical analysis, performed using Wilcoxon signed rank test (to test whether CI is over 1) \**P* = 0.0142. G. Proposed model of *E. coli* growth upon confinement. At the single-cell scale, *E. coli* adaptation to mechanical stress is described in 3 phases. In a first phase, bacteria face a lack of space limiting their growth (magenta), thereby uncoupling growth and protein synthesis. Consequently, protein concentrations increase (green), leading to a first increase in cytoplasmic crowding (dark red) sufficient to trigger bacterial division (black). In the meantime, bacteria activate the Rcs envelope stress response. In a second phase, the uncoupling between growth and division leads to the formation of tiny bacteria characterized by a higher DNA occupancy (blue). This results in an additional increase in crowding, which further inhibits bacterial division. In a third phase, while growth, division and protein synthesis are arrested, bacteria continue increasing their turgor pressure and require Rcs-mediated envelope remodeling (red) to maintain their shape upon *overpressurization*. At the global scale, this increase in turgor pressure results in the generation of a large growth-induced pressure onto the bacterial microenvironment (orange). Time is indicated as hh:mm. All scale bars: 5 µm.

To explore the relevance of these results to infection, we asked whether the Rcs pathway was also involved in bacterial adaptation to confinement during UTIs *in vivo*. To test this, we deleted *rcsB* using two different approaches and fluorescent labels (GFP and mKate) in the UPEC strain UTI89 and tested the phenotype in the bacterial confiner (**Figure 6.E**). We observed that the *rcsB* mutant in the pathogenic strain also resulted in large amorphous cells. Synthesis of colanic acid capsule played a partial role in shape maintenance as the mutant *wcaJ* displayed a subtle morphological phenotype (**Supp Figure 7.I-L, Supp. Video 7**). To determine whether the loss of Rcs pathway might impact UPEC fitness and virulence *in vivo*, mice were intravesically infected with an equal mix of UTI89 wild-type and *rcsB* mutant bacteria. The competitive index, calculated as the number of *rcsB* divided by wild-type cells recovered from the bladder 24 hours post-infection was less than 1, indicating that the loss of Rcs led to a decreased fitness for UPEC *in vivo* (**Figure 6.F, Supp. Figure 7. M, N**). Altogether, these results show that the Rcs pathway is essential to maintain bacterial shape in the *overpressurization* regime, and contributes to the fitness of uropathogenic *E. coli*, suggesting that it might constitute an interesting target for therapy.

## DISCUSSION

Using a microfluidic device adapted to the study of confined growth of bacterial colonies, we identified a regime of stalled growth, division arrest and repressed protein synthesis during which turgor pressure steadily increased, reaching hundreds of kPa, that we name the *overpressurization* regime. We found that the Rcs pathway is activated in this regime and that this activation is required for mechanical adaptation and for fitness of pathogenic *E. coli* in a mouse model of urinary infection. Thanks to quantitative approaches and physical modeling, we were able to propose a simple mechanistic working model for confined bacterial growth, leading to the *overpressurization* regime (**Figure 6.G**): (*i*) In the absence of spatial limitation, cell growth and protein and osmolyte synthesis are tightly coupled to regulate cell division and maintain size homeostasis; (*ii*) When space becomes limited, bacterial proliferation induces pressure build-up; (*iii*) During the first 30 minutes post-confinement, bacterial growth rapidly declines, inducing an increase in protein concentrations, which triggers cell division. This leads to a rapid increase in cell number while cell size decreases (Phase 1); (*iv*) From 30 minutes to 2 hours post-confinement, protein accumulation and increased DNA occupancy induce an increase in cytoplasmic crowding, arrest in protein synthesis, and a sharp decrease in cell division, while pressure increases steadily (Phase 2); (*v*) After 2 hours, protein synthesis is arrested, but the pressure produced by bacteria still increases in a linear fashion. Based on our model, we propose that this is due to a continuous accumulation of trapped osmolytes whose accumulation in cells does not depend on protein synthesis but on protein activity (Phase 3). In this last *overpressurization* regime, activation of the Rcs pathway is required to maintain cell shape by strengthening the cell envelope in order to sustain the increasing pressure.

During the different confinement phases, bacteria steadily build-up a large compressive pressure, reaching up to 300 kPa, in agreement with recent measurements of turgor pressure in individual cells(*48*, *49*) and in biofilm growth on adhesive substrates(*9*, *50*). Confinement of eukaryotic cells also generates growth-induced pressure, with magnitudes scaling with osmotic pressures exerted on the cell wall in the case of yeast and on the actin cortex in mammal cells(*12*, *14*, *15*). These results reinforce the idea that the main source of confinement-induced mechanical stress produced by a multicellular colony is the osmotic pressure generated by single cells.

While in homeostatic conditions growth and division are coupled to maintain a constant cell size, in the first phase of confined growth, the two get uncoupled, leading to major changes in cell volume. Pressure build-up induced a sharp slow-down in bacterial growth. In the meantime, bacteria rapidly undergo multiple cycles of reductive divisions to finally arrest at a minimal cell size. Rod shortening was reminiscent of stationary phase cells, however in our confinement device fresh medium was constantly perfused, preventing starvation(*51*, *52*).

While the impact of confinement on growth rate appears general to all living systems explored so far(*12*, *53*, *54*), its effects on the cell division cycle are very diverse. Budding yeast maintain a constant cell volume under mechanical pressure, by arresting division when growth slows down(*13*). Epithelial cells, similarly to our observations in bacteria, display an uncoupling between growth and division rate in dense monolayers, which also induces a global reduction of cell volume, reaching a minimal value defined by genome size(*15*, *29*). While the increase in karyoplasmic ratio is detrimental in the epithelium leading to DNA damage, we showed that *E. coli* has an extreme plasticity to changes in DNA occupancy and rapidly returned to their normal size and division cycle upon pressure release without DNA damage(*29*, *55*). These results point out to the remarkable adaptation of bacteria to growth under pressure.

We found that one key feature of bacterial confinement at short timescales is an increase in intracellular molecular crowding, occurring primarily in Phases 1 and 2. Cytoplasmic crowding influences the activity of proteins and macromolecular complexes in all cells, including bacteria(*56*, *57*). We showed that at early stages of confinement, protein concentrations increased and this process, reproduced by osmotic shocks, was sufficient to induce an uncoupling between division and growth. This is explained by the increased concentration of divisome machinery proteins such as FtsZ and rules out a direct role of turgor pressure(*34*, *35*). Later on, increase of DNA occupancy mediates transition of the cytoplasm to a state similar to the previously described “colloidal glassy” state upon ATP or nutrient depletion(*58*), but in our case triggered by a mechanical cue.

An apparent paradox in our results concerned the increase in growth-induced pressure at late stages of confinement (Phase 3) while protein production is already arrested, raising the question of the origin of this sustained increase in pressure. We propose that while proteins cease to increase in concentration, enzymes remain active leading to metabolite accumulation and turgor increase. This idea is supported by the minimal theoretical framework developed in this study and points to the mechano-osmotic role of metabolism. Unlike proteins whose role on growth has been the focus of several studies in the past few years, the osmotic contribution of metabolites such as glutamate to cell volume is a new concept in the field and an emerging mechanism of cell size control(*38–40*, *42*). Further studies are needed to understand how synthesis of small metabolites is impacted by cytoplasmic crowding and regulates growth-induced pressure in the presence of confinement(*59*).

A major prediction of the theoretical model is that confined bacteria, if they are constantly provided with nutrients and thus able to accumulate trapped osmolytes, will undergo a massive internal turgor pressure increase while maintaining a proper morphology. We found that this activates stress responses, with a crucial role of the Rcs pathway in bacterial adaptation to mechanical confinement(*20*, *46*, *60*). Strikingly, in the absence of the *rcsB*-dependent transcriptional response, *overpressurized* (Phase 3) cells display large and heterogeneous shapes reminiscent of L-forms(*61*, *62*). The mild phenotype observed in the capsule-deficient mutant *wcaJ* is consistent with previous findings suggesting contribution of this structure to global cell mechanics(*20*) (**Supp. Figure 7**). The transcriptional regulation on other components of the bacterial envelope by the Rcs pathway, such as peptidoglycan organization and the outer membrane, are likely main contributors to cell shape maintenance in the *overpressurization* regime(*18*, *45*, *63*). Interestingly, we found that the Cpx stress response is also transcriptionally activated during confined growth, but the corresponding mutant did not display any major shape defect(*44*, *64*). The Rcs pathway is therefore central and specific to bacterial adaptation to turgor pressure increase during confinement. An interesting aspect of this global transcriptional reprogramming that we have not investigated in detail in this study is that the spatial pattern of activation is not homogeneous throughout the chambers and is specific to each stress response pathway. Further studies with our confinement device will allow a deeper understanding of the complex interplay between different stress responses and how integration of mechanical signals coordinates multicellular growth and cell fate specification in bacterial biofilms, as recently shown during organogenesis(*16*). It might also reveal other pathways required for bacterial survival during confined growth.

Importantly, uropathogenic *E. coli* also underwent rod shortening during confined growth, a feature previously described during bladder infection(*23*). This suggests that these bacteria are experiencing mechanical confinement. Accordingly, an *rcs*-deficient UPEC strain showed decreased fitness during infection of the uroepithelium *in vivo*. Previous studies showed that intracellular UPEC growth was associated with lower antibiotic susceptibility(*65*) and the Rcs pathway is also known to promote antibiotic tolerance(*66*). Further studies are needed to determine whether confinement is sufficient to increase bacterial survival to antibiotic treatment, and whether this is dependent on the Rcs pathway activation. More generally, we anticipate that our experimental approach and physical model might help to decipher bacterial virulence mechanisms associated to mechanical constraints.

In conclusion, this work uncovers novel molecular and physical mechanisms responsible for bacterial adaptation to mechanical constraints with important implications for bacterial evolution, antibiotic resistance and bacterial infections.

## Acknowledgements

The authors thank C. Beloin, I. Matic, M.-F. Bredeche, M. Micaletto, J. Bos, T. Bernhardt and A. Lindner for providing bacterial strains and helping with the cloning strategy; K. Kline for suggesting to us to work with pathogenic strains; L. Deltourbe for performing *in vitro* experiments of infections; S. Gobaa, J. Wong, A. Salles and members of the Biomaterials & Microfluidics and Photonic BioImaging facilities at Institut Pasteur; A. Lecestre and all the engineers of the LAAS-CNRS clean room for technical assistance; M.-G. Côme for providing built-in scripts for image acquisition; O. Disson in M. Lecuit’s lab for giving access to a microscope equipped with a CellASIC system; P. Nivoit for editing all supplementary videos present in the paper; C. Beloin and I. Matic for fruitful discussions and feedbacks; S. Rigaud from the Image Analysis Hub at the Institut Pasteur for initial guidance in image analysis; A. Persat, P. Sens, B. Baum, C. Beloin, I. Matic for reading the manuscript.

## Author contributions

LLB, MD, GD, and DB designed the study. LLB designed, performed and analyzed most experiments and actively contributed to manuscript preparation by building figures and writing the manuscript. BA, LLB and MD designed the bacterial confiner and optimized the protocol for microfabrication. RR built up a theoretical model of bacterial growth under confinement and edited the manuscript. LX and LLB built up the pipelines for bacterial segmentation, tracking and post-processing and performed image analysis of the microscopy datasets. LRF performed *in vivo* and *ex vivo* urinary tract infections in mice. SG constructed most of the bacterial strains used in this study, together with LLB. LM provided crucial help with microfabrication of the nanochannels in the confiner. MAI supervised LRF during the infection experiments *ex*/*in vivo*, helped write and edit several parts of the manuscript, and provided expertise on UPEC and UTI. MP gave essential conceptual feedback on the theoretical model, suggested key experiments, provided feedback throughout the study and helped write and edit several parts of the manuscript. JYT supervised LX, provided feedback on all the image analysis present in this study and helped with study conceptualization. MD, GD and DB developed the initial hypothesis, supervised the study and wrote the manuscript. All authors contributed to manuscript preparation.

## Data and code availability

Any additional information required to reanalyze the data reported in this paper is available from the lead contact upon request. All the data and codes required to reproduce the main Figures are publicly available on Zenodo as of the date of publication (https://zenodo.org/records/12799692). DOIs are listed in the key resources table.

