## Supplementary Materials for "Bacterial growth under confinement requires transcriptional adaptation to resist metabolite-induced turgor pressure build-up"

#### Materials and Methods

##### Bacterial strains

The bacterial strains, plasmids and primers used in this study are described in **Tables 1 and 2**. Genetically modified *E. coli* strains were derivatives of the following *E. coli* strains: the K-12 MG1655, TB28, BW27783 strains, one from the XL1-Blue background and the uropathogenic cystitis isolate UTI89 strain(*30*). Bacteria were grown in liquid or solid Luria-Bertani medium (LB, BD Difco) at 37°C or 30°C when grown to perform allelic exchange experiments. Antibiotics were used at the following concentrations: ampicillin (Ap), 100 µg/ml, chloramphenicol (Cm), 25 µg/ml, kanamycin (Km), 25-75 µg/ml, tetracycline (Tc), 2.5 µg/ml. Genetically modified *E. coli* strains were constructed either by λ-red recombination or P1 phage transduction, as described in the following.

For λ-red recombination, PCR products (detailed below) were transferred by λ-red recombination method into *E. coli* K-12 MG1655 and derivatives or UTI89 and derivatives transformed with the pKOBEGA or pKOBEG plasmids(*67*), which contains the λ-Red operon under the control of the arabinose-inducible araBAD promoter. PCR fragments were dialyzed on a 0.025 µm porosity filter (Millipore) and electroporated after induction with 0.2% arabinose. Bacteria were spread on LB agar plates with the appropriate selection antibiotic and incubated overnight at 37°C. Transformants were checked by PCR and sequencing and then tested for ampicillin or chloramphenicol resistance to test loss of pKOBEGA or pKOBEG, respectively.

The PCR products that have been generated are described below:

- ZipA(TM)-mCherry2-Tc$^R$: A fragment containing ZipA(TM)-mCherry2 and a Tc$^R$ cassette was created in two steps. First, the Tc$^R$ cassette was amplified from pBR322 vector(*68*) with primers HK022-att-Tc-F/ Tc-R, and the ZipA(TM)-mCherry2 fusion was amplified from *E. coli* TB28(attHKpHC503)(*26*) with Tc-PzapA-F/HK022-att-mCherry-R. The latter PCR product plus the amplified Tc$^R$ cassette were used for a two-way overlap PCR with primers HK022-att-Tc-F and HK022-att-mCherry-R. The *tet* gene was oriented in the opposite direction to ZipA(TM)-mCherry2. Primers HK022-att-Tc-F and HK022-att-mCherry-R carried 50 bp of homology with the insertion region at each end (HK022 att site). The expected chromosome structure in the HK022 att site was confirmed with primers P1 and P4 (**Table 2**)(*69*). The PCR product was inserted into MG1655, MG1655 GFP, MG1655 HU-GFP, MG1655 $P_{cpxP}$-meGFP, MG1655 $P_{recA}$-YFP, MG1655 GEM40 and BW27783 FtsZ-mNG.

- GEM40-Cm$^R$ : The sequence encoded for the 40 nm GEM was amplified using GEM40-XhoI-F/GEM40-PmeI-R primers from the plasmid pCDNA3.1-pCMVPfV-GS-Sapphire (Addgene #116933)(*33*) and cloned into pMGC10 vector(*28*) with XhoI and PmeI restriction sites. The plasmid contained the open reading frame encoding the *Pyrococcus furiosus* encapsuling 196 fused with the T-Sapphire fluorophore under the control of the inducible P*lac* promoter. Then a fragment 40 nm GEM containing a Cm$^R$ cassette was created in two steps. First, the Cm$^R$ gene was PCR amplified from *E. coli* TG1_λATT-Cm-GFP (UGB1479(*70*)), and the 40 nm GEM was amplified from the previously constructed pMGC10 plasmid with Cm-UGB1479-F_λATT-Sapphire-R. The latter PCR product plus the amplified Cm$^R$ cassette were used for a two-way overlap PCR with λATT-Cm-F/λATT-Sapphire-R. The *cat* gene was oriented in the opposite direction to 40 nm GEM. Primers λATT-Cm-F/λATT-Sapphire-R carried 41 bp of homology with the insertion region at each end (λATT site). The expected chromosome structure in the λATT site was confirmed with primers λATT-ext5-F and λATT-ext3-R. The PCR product was electroporated into both XL1-Blue and MG1655 backgrounds. In the latter, the low-copy plasmid pREP4 (Qiagen) was transferred into *E. coli* MG1655 ZipA-mCherry strain expressing GEM40, constitutively expressing the LacI repressor protein and tightly regulating recombinant protein.

- HU-GFP, *slmA, wcaJ, cpxR, rcsB*: PCR fragments containing a $Km^R$ cassette were amplified from *E. coli* SJ156(*32*) or from the Keio collection(*71*) respectively with primers listed in **Table 2**.

- mKate: a PCR fragment containing a $Cm^R$ cassette was amplified from *E. coli* MG1655 mKate(*72*) with primers mKate-F and mKate-R **(Table 2)**

- P1 allelic transduction from the Keio collection(*71*) was used to delete the *rcsB* gene into *E. coli* MG1655 ZipA-mCherry. Transductants were verified by PCR and sequencing, with the following primers listed in **Table 2**.

**Bacterial culture**

Bacteria were streaked from -80°C frozen stocks onto fresh LB agar plates supplemented with the appropriate antibiotics and incubated overnight at 37°C. For the experiments carried out with the MG1655 strain, a few bacterial colonies from plates were resuspended in 5ml filtered LB Miller and incubated for 7-12 hours at 37°C with agitation at 180 RPM. A bacterial subculture was then started in pre-warmed filtered LB Miller at an optical density of 0.01 and incubated for 2 hours at 37°C with agitation at 180 RPM. For the experiments carried out with the UTI89 strain, a few bacterial colonies from plate were resuspended in 20ml filtered LB Miller and incubated for 12 hours at 37°C without agitation as is standard for bacterial preparations used for *in vivo* infection. For *ex vivo* and *in vivo* experiments, a single bacterial colony from a plate was resuspended in 10ml LB Miller supplemented with the appropriate antibiotics and incubated for 18 hours at 37°C without agitation to allow stationary cells to express type 1 fimbriae.

**Bacterial stress reporter validation**

Bacterial precultures were diluted at an optical density of 0.1, loaded in 96 well black-bottom plates, submitted to a range of chemical stress inducers, and incubated in the plate reader (Cytation5, Biotek) under agitation at 37°C with automated acquisition of optical density and fluorescence every 30 minutes for 18 hours. For the chemical inducers, cephalexin and ciprofloxacin were used to induce envelope and DNA damage respectively. Data were analyzed on GraphPad Prism software.

**Bacterial proliferation on agar pads**

Bacterial proliferation on agar pads was used to assess bacterial growth in classical conditions. For this, exponentially growing bacteria diluted at an optical density 0.015 were trapped between a glass coverslip and a 2% agarose gel composed of LB Miller, placed under the microscope, and imaged at 37°C (The Cube).

**Microfabrication**

Masters containing the chip design were fabricated in the clean room at the LAAS-CNRS using two-layer stepper-based high-resolution photolithography. The designs of the silicon master molds were created using Clewin software by rescaling the dimensions of the chip previously developed to confine yeasts(*12*). Briefly, the first layer defining the nutrient channels was patterned on a 1.1µm layer of ECI positive photoresist using a **UV projection stepper** (**Canon FPA 3000i4/i5**, dose = 400J/m$^2$) and etched at a depth of 400nm by *high density plasma etching*. The second layer defining the growth chambers and the loading channels was patterned on SU8 negative photoresist at a height of 2.6µm. For this, the silicon wafer previously etched was coated with SU8 negative photoresist, aligned with the first layer, and insolated in the stepper using a dose of 3000J/m$^2$. Characterization by profilometer and scanning electron microscope was performed between each step to ensure a correct aspect ratio. Before being taken out of the clean room, the masters were passivated with silane. For chip preparation, a

15:1 mixture of PDMS (Sylgard 184, Dow corning) and curing agent was deposited in drops of roughly 100µl on top of the chamber features and cured at 65° for 30 minutes. A 10:1 mixture was then poured and cured overnight at 65°C. Then, PDMS was cut, punched, cleaned with isopropanol, bounded to pre-cleaned glass coverslips (Marienfeld, #1.5) using oxygen-plasma (Femto Science, power = 60W, exposure time = 40s), cured at 65°C for 5h, sterilized using UV ozone cleaner (UVO Cleaner, Jelight) for 30 minutes and finally cured at 65°C for 30 minutes. Prior to all experiments, the integrity of the nanochannels was checked.

**Microfluidic device operation**
The microfluidic chips were filled in the following order to prevent air bubbles within the channels(*13*) using sterile equipment. First, exponentially growing bacteria were loaded at an optical density of 0.3-0.4 in the growth chambers. Second, the loading channel was filled with pre-warmed filtered LB Miller using a Fluigent pressure controller (MFCS, Fluigent) set at 1 bar during the whole course of the experiments. Third, a plug was inserted at the outlet of the loading channel to allow continuous medium renewal within the growth chambers. The chips were then placed in the microscope thermoregulated chamber set at 37°C.

**Microfluidic device calibration for pressure measurements**
Growth-induced pressure was inferred from the deformation of the chambers as previously done(*73*). To optimize the detection of the chamber walls, the rigidity of the PDMS was adjusted and the channel edges were fluorescently stained by perfusing LB Miller supplemented with FM1-43 at 2µg/ml. Chamber deformation was semi-automatically tracked using homemade Fiji and Python scripts. The relationship between deformation and growth-induced pressure was then calibrated as follow: the deformation of the loading channel was measured at the top of the channel under a range of known pressures (0, 200, 400, 600, 800, 1000, 1500, 2000, 3000, 4000mbar). The applied pressure was represented as a function of the measured deformation and fitted with a linear regression curve ($y = 282.6x$, $R^2 = 0.99$) (**Supp. Figure 1.G**). This calibration curve was established for all the chips where force measurements were performed. To facilitate force measurements while looking at other fluorescent reporters, we also calibrated the growth-induced pressure as a function of the number of bacteria in the 2D focal plane of observation. This provided a strategy to infer the pressure in the chamber by using the number of bacteria as a readout (**Supp. Figure 1.I**).

**Osmotic shocks**
Hyperosmotic shocks were performed by increasing the osmolarity of the culture medium using sorbitol (Sigma-Aldrich) at various concentrations (0.25, 0.5, 1M) as previously done(*18*, *74*), either using Ibidi channels or CellASIC ONIX commercial chambers when continuous flow perfusion was required.
In the first assay, exponentially growing bacteria were introduced slowly within Ibidi channels (sticky-slide VI 0.4, Ibidi) coated with 3-Aminopropyltriethoxysilane (APTES, Sigma-Aldrich) at 2% in ethanol and incubated for 10 minutes at 37°C to promote bacteria attachment. LB medium was then rinsed out and replaced with fresh medium supplemented with sorbitol at various concentrations (0.25, 0.5, 0.75, and 1M). Image acquisition was launched 30 seconds after changing the medium. Note that *E. coli* XL1Blue bacteria were used for these experiments.
In the second assay, exponentially growing *E. coli* bacteria fluorescently labeled at the inner membrane and DNA were loaded in B04A microfluidic perfusion plates (CellASIC) and pre-warmed medium was exchanged using the ONIX microfluidic platform (Cell ASIC). The culture medium was switched with culture medium supplemented with sorbitol at 0.5M first, and then 1M to progressively increase cytoplasmic crowding. The culture medium was always supplemented with wheat germ agglutinin-AlexaFluor 642 (WGA-AF642, Invitrogen) to

fluorescently label the cell wall. Bacterial division was monitored based on the inner membrane signal and confirmed with cell wall fluorescent staining in a time window of 3 minutes after the hyperosmotic shock as previously done(*35*). For quantification, bacteria with a clear membrane invagination and delimitation at mid-cell were considered as dividing. The fraction of dividing cells was determined by manually counting the number of dividing bacteria over the total number of bacteria.

**Fluorescent probes**
Several fluorescent probes were perfused in the bacterial confiner to characterize bacterial aggregates upon confinement. SYTOX Green (Invitrogen) was supplemented in the culture medium at 5 µg/ml to fluorescently stain nonviable bacteria characterized by a permeable envelope. FITC-Dextran 10kDa (Sigma-Aldrich) was supplemented in the culture medium at 1 mg/ml to assess the kinetics of medium perfusion within confined aggregates. The amphiphilic dye FM1-43 (AAT Bioquest) was supplemented in the culture medium at 2 µg/ml to fluorescently stain PDMS walls.

**Live imaging**
Most of the experiments were performed using a spinning-disk confocal microscope (Ti Eclipse, Nikon) equipped with a spinning-disk module (CSU-X1, Yokogawa) and a 100X oil objective (Plan Fluor, NA = 1.45) coupled to a Live-SR SIM-like module (Gataca Systems). The microscope stage was kept at 37°C using a temperature controller (The Cube). Fluorescence was recorded using a CMOS camera (95B Prime, Photometrics). The laser power was set at 30%, the exposure time was set at 300ms, and the focus was maintained using a Perfect Focus System (PFS, Nikon). Image acquisitions were computer-controlled using MetaMorph software and custom-made scripts (Molecular Devices). The CellASIC experiments (**Figure 4.B**) were imaged using a confocal microscope (LSM710, Zeiss) equipped with an Airyscan module (Zeiss) and a 63X oil objective (Zeiss, NA = 1.4). Super-resolution 3D imaging of the chips (**Supp. Figure 2.A**) was performed at a fixed timepoint using a SIM microscope (Zeiss Elyra7, Lattice SIM) equipped with a 63X oil objective (Plan Apochromat, NA = 1.4 DIC M27). Similar Z-stacks acquisitions were performed using 1µm-wide beads to correct optical aberrations (Tetraspeck, Invitrogen).
Unless stated, bacterial proliferation in the bacterial confiner was monitored with a timelapse of 30 minutes frame rate over 12-17 hours and Z-stacks of 0.1 µm step over 1µm centered around the focal plane corresponding to the nanochannels. For force measurement, a MetaMorph script was used to acquire Z-stacks of 0.2 µm over 2.6 µm to monitor chamber walls at the height of maximal deformation. For tracking analyses, bacterial proliferation was monitored at 5 minutes frame rate for 5-6 hours only to avoid photobleaching. Bacterial proliferation in agar pads was monitored with a timelapse of 5 minutes frame rate over 2.5 hours to avoid nutrient depletion. GEM40 diffusion was imaged in stream mode every 50 milliseconds over 2.5 seconds using a laser power set at 95%.

**Mouse model**
Mice had free access to water and food and experiments were carried out under the protocol APAFIS #34290, approved by SC3–CEEA34–Université de Paris Cité, at Institut Cochin, in application of the European Directive 2010/63 EU. Mice were anesthetized with ketamine (100 mg/kg) and xylazine (5 mg/kg) intraperitoneally and sacrificed by cervical dislocation after isofluorane anesthesia.

***Ex vivo urinary tract infection***

Bladders from naïve 12-week-old male C57BL6/J mice were provided from a team at Institut Cochin (implementing the 3R method of reducing mice number). Bladders were cut in quarters and the epithelium was peeled away from the muscle layer. Urothelium quarters were infected with UPEC for 24 hours at 37°C. Urothelial quarters were washed in 1X PBS twice to remove loosely attached bacteria from the surface. The quarters were fixed in 4% PFA for one hour at 4°C (when infected with UTI89-RFP) or pinned on 2% agarose and then fixed in 4% PFA for one hour at 4°C (with mKate or GFP strains). Quarters were blocked and permeabilized (1% bovine serum albumin, 0.3% triton X-100, 1% donkey serum) overnight at 4°C while shaking. Quarters were washed, stained with DAPI (1:1000) for one hour at 4°C while shaking. Urothelium sheets were clarified with RAPI clear 1.47 for 3 days minimum and images were acquired using a spinning-disk confocal microscope (Ti Eclipse, Nikon) equipped with a spinning-disk module (CSU-X1, Yokogawa) and a 100X oil objective (Plan Fluor, NA = 1.45) coupled to a Live-SR SIM-like module (Gataca Systems).

***In vivo* urinary tract infections**

Female C57Bl/6 mice, aged 6-7 weeks (Charles River Laboratories, France) were intravesically infected with $10^7$ CFU/ml of an equal mix of wildtype and *rcsB* mutant strains (of the same background) in 50 µl of sterile 1X PBS(*75*, *76*). Mice were sacrificed 24 hours post-infection and bladders were removed and homogenized in 1ml of sterile 1X PBS. Bladder homogenates were plated on selective agar plates containing the appropriate antibiotics to count wildtype and *rcsB* mutant strains. The values of competitive indexes were calculated as colony-forming units of *rcsB* mutant strain over colony-forming units of wildtype strain(*77*).

***Image analysis***

***All the analysis scripts and example data to reproduce the analysis are available in the*** Supplemental Material***. Briefly:***

*Force measurements*

Fluorescent images of chamber walls at a height of 2.6µm (*i.e.* the top of the chamber at the beginning of the experiment – but also the height at which the chamber is the most deformed during the whole course of the experiment) were binarized using the Otsu auto-threshold in Fiji and verified manually. The distance along the x-axis between the chamber walls was measured using a Python script and used to compute the deformation over time with respect to its initial size. Similar analysis was performed on the calibration channel to establish the calibration curve. The deformation of the chamber was then related to the corresponding growth-induced pressure by using the equation established during the calibration.

*Bacterial single-cell segmentation and tracking*

Single bacteria were segmented using ilastik(*78*) (v1.3.3) pixel-classification workflow, trained specifically on the bacterial inner membrane fluorescent signal. The model file generated from manual annotations was then used in TrackMate, using its ilastik integration(*28*) (v7.11.1) to segment bacteria with a threshold on the probability map of 0.6. For the morphology measurements reported in **Figure 2**, segmentation mistakes were manually corrected using Fiji on the exported mask images. To measure bacterial division and growth rates (**Figure 2**), the segmented bacteria were tracked with TrackMate using the overlap tracker with parameters: min IoU 0.1, scale 1.3. Tracking mistakes were manually corrected in TrackMate.

*Karyoplasmic ratio*

The DNA occupancy in single bacteria was determined by segmenting the DNA channel in images (**Supp. Figure 3.C**). As above, we used ilastik pixel classification workflow, trained a model on the DNA channel further used in TrackMate with a threshold on the probability map of 0.7. The Python script to measure the karyoplasmic ratio from the bacterial and DNA masks is given in the Supplemental Material.

*Single bacteria measurements*

All the bacterial segmentation data generated with TrackMate and ilastik, with or without tracking data, were analyzed subsequently with dedicated Python scripts, tailored for the specific feature to extract. Fluorescence intensity measurements and bacterial area were extracted from the numerical features in TrackMate objects. Bacteria width and length were measured following the method described below.

*Bacterial width and length measurements*

Bacterial width and length were measured using a custom approach in Python (**Supp. Figure 3.C**). The skeleton of each bacterium was generated from the contours stored in the TrackMate files, then pruned and smoothed to obtain the bacterium medial line. The bacterial length was defined as the length of the medial line, plus the distance from the medial line extremities to the border, measured with the distance map. The bacterial width was defined as the median of the width distribution measured along the medial line, again using the distance map. The shape irregularity index was measured as the standard deviation of this distribution normalized by its mean value for the wild-type strain (**Figure 6**).

*Protein production rate*

The protein production rate (PPR) of a fluorescent protein was measured over time as the variation of the protein total fluorescent intensity (TFI) over the whole chamber divided by the number of bacteria N, i.e. PPR = $\frac{1}{N} * [\frac{1}{TFI}\frac{d(TFI)}{dt}]$. This measurement cannot be done at the single-cell scale because of the volume changes induced by the reductive divisions.

*Determination of transcriptional reporter activation*

Bacteria expressing fluorescent transcriptional reporters ($P_{rcsA}$-GFP, $P_{cpxP}$-meGFP, $P_{recA}$-YFP) were classified as activated or non-activated based on a threshold on the fold-change of fluorescence intensity, with respect to movies frames where bacteria were not confined. This threshold was manually determined, based on the shape of the histogram of the fold change distribution of intensity of all movies (**Supp. Figure 7.C-E**).

*Spatial pattern of Rcs transcriptional response*

In **Figure 6.A**, the contour delineating a zone of lower Rcs activation in the chambers was drawn manually to roughly include all bacteria for which RcsA intensity was below a threshold. This threshold was determined by the Otsu method on the distribution of RcsA intensity measured on all bacteria in 3 chambers. Then, bacteria were classified as 'Center' or 'Edges' depending on whether they were found respectively inside or outside this contour (**Figure 6.D**).

*Ex-vivo infection image analysis*

The 3D stacks obtained from imaging bacteria in the *ex vivo* assay were manually segmented in Fiji as described in the following. A single Z-slice was selected in the middle of the volume and manually delineated in 2D. Ambiguities were resolved by inspecting Z-slices directly above or below the selected one. The resulting label image was imported in TrackMate and analyzed as described above.

*40nm-GEMs diffusion measurements*

Single 40nm-GEMs nanoparticles were tracked using the MOSAIC Fiji plugin(*79*) using the following parameters: size = 3 pixels, cutoff = 0, link = 1 frame, displacement = 5 pixels per frame. Single-cell tracks longer than 10 points were analyzed to compute the corresponding mean squared displacement (written MSD) and diffusion coefficient using a MATLAB code developed in a previous study(*33*).

**Data visualization and statistical analysis**

Number of replicates and number of tracks / bacteria / chambers / aggregates analyzed are specified in corresponding legends. All the observations described in the bacterial confiner were highly reproducible and observed in more than 15 chambers. Complex single-cell analyses were completed on a representative subset of data ($n_{chambers}$ = 2 - 4) for the sake of time. Quantifications still consider a large number of individual bacteria ($3928 \leq n_{bacteria} \leq 50661$). Details on statistical analyses are written in figure legends. For all the experiments, data visualization and statistical analysis were performed on GraphPad Prism.

### Supplementary Model: "Bacterial growth under mechanical confinement requires Rcs pathway activation to sustain overpressurization"

Laure Le Blanc, Baptiste Alric, Romain Rollin, Laura Xénard, Laura Ramirez-Finn, Sylvie Goussard, Laurent Mazenq, Molly Ingersoll, Matthieu Piel, Jean-Yves Tinevez, Morgan Delarue, Guillaume Duménil and Daria Bonazzi

#### I. MODEL

In the following, we derive a model to understand quantitatively the growth of a bacterial colony under confinement.

##### A. Evolution of the GIP in the chamber

In recent years, more and more evidence has pointed to the key role of osmosis on cellular and organelle growth ([1],[2], [3], [4]). We take into account this physical principle in our model by imposing that the chemical potential of water is balanced throughout the bacterial membrane. This is equivalent to enforcing the equality between the difference in osmotic pressure with the difference in total mechanical pressure applied throughout the bacterial membrane ([1]). Here, at the difference of single-cell growth, the total mechanical pressure is composed not only of the hydrostatic pressure $P$ but also of the external mechanical stresses $\sigma^{el}$ due to the bacterial crowding in the chamber.

$$\Pi^b - \Pi_0 = \Delta P + \sigma^{el} \tag{1}$$

Where, $\Pi$ accounts for the osmotic pressure within the bacteria, subscript $_c$, or in the external medium, subscript $_0$; $\Delta P$ is the difference of hydrostatic pressure throughout the bacterial membrane. It is related to the tension $\gamma$ and the average mean curvature $\frac{1}{w}$ of the cell wall, through the Laplace law $\Delta P = \frac{4\gamma}{w}$ (see figure). For the sake of simplicity, we assume that cell wall constituents are incorporated during bacterial growth to maintain $\Delta P$ ([5]). Moreover, as most of the osmolytes within bacteria are accounted for by small osmolytes, we consider that the osmotic pressure is dominated by the ideal gas term:

$$\Pi^b = k_B T \cdot \frac{N^b}{V^b - R^b} \tag{2}$$

Where, $N^b$ is the number of osmolytes trapped inside the bacteria, $V^b$ is the volume within the bacteria, and $R^b$ is an excluded volume that depends on the macromolecular content in the bacteria. As most of the dry content of the bacteria is accounted for by proteins, we will assume that this excluded volume is simply proportional to the number of proteins within the bacteria $P^b$ times an average effective volume occupied per protein $v_p$, $R^b \sim v_p \cdot P^b$.

For simplicity, we will also consider average behaviors within the bacterial colony. We thus relate, the quantities at the bacterial scale, upper script $^b$, and at the colony scale, no upper script, by assuming that the bacteria are confluent in the plan of observation (bottom of the chamber of fixed total area $A_c$, Fig.5.A).

$$A_c = n^{2D} \cdot \left(A^{b,proj} + A^{\mathcal{I}}\right) \tag{3}$$

Where $n^{2D}$ accounts for the number of bacteria in the plan of observation, at the bottom of the chamber. We also assumed that the space in the chamber was divided into two contributions. (1) The bacterial protoplasms of average projected area $A^{b,proj}$. (2) And an interstitial space of projected area $A^{\mathcal{I}}$, which experimentally corresponds to the fluorescent delimitation between two bacteria. We further approximate the bacteria as a cylinder of long axis length $l$ and diameter $w$ enclosed by a shell of width $\delta$ (Fig.5.B). Note that the previous geometrical constraint is true up to a constant coefficient that won't change our conclusions. Moreover, we verify that this approximation is in good agreement with our experimental quantifications (Fig.1.A). Based on these geometrical considerations we further express the average protoplasm volume and projected area as:

$$V^b = \frac{\pi}{4} \cdot w \cdot A^{b,proj} \quad \text{with,} \quad A^{b,proj} = \left(\frac{A_c}{n^{2D}} - 2 \cdot \overline{\delta} \cdot w^2 \cdot (1 + 2 \cdot \overline{\delta})\right) \cdot \frac{1}{1 + 2 \cdot \overline{\delta}} \tag{4}$$

With, $\overline{\delta} = \frac{\delta}{w}$. We finally impose that the density of bacteria $\rho$ in the chamber is homogeneous. This allows us to relate the total number of bacteria $n$ in the volume $V$ of the chamber with the number of bacteria in the plan of observation as:

$$\rho = \frac{n}{V} = \frac{n^{2D}}{A_c \cdot w} \tag{5}$$

Combining Eq.5 and Eq.4, we finally express the bacterial protoplasm average volume $V^b$ as:

$$V^b = \left(\frac{V}{n} - 2 \cdot \overline{\delta} \cdot w^3 \cdot (1 + 2 \cdot \overline{\delta})\right) \cdot \frac{\pi/4}{\left(1 + 2 \cdot \overline{\delta}\right)} \tag{6}$$

Furthermore, mechanical stress builds up in the chamber due to the chamber crowding. We call this stress the growth-induced pressure -GIP - and we relate it to the volume of the chamber $V$ through a scalar Hooke's law:

$$\sigma^{el} = E \cdot \left(\frac{V}{V_c} - 1\right) \tag{7}$$

Where, $E$ is the elastic modulus of the chamber and $V_c$ is the non-deformed volume of the chamber. Using Eq.2, Eq.8, and Eq.7 we show that the volume of the chamber is the solution of a second-order polynomial that reads:

$$\overline{V} = -\frac{1}{2} \cdot \left((\overline{\Pi}_0 - 1) \cdot \overline{V}_c - \overline{V}^e \cdot \overline{P}\right) + \frac{1}{2} \cdot \sqrt{\left((\overline{\Pi}_0 - 1) \cdot \overline{V}_c + \overline{V}^e \cdot \overline{P}\right)^2 + 4 \cdot \left((\overline{\Pi}_0 - 1) \cdot \overline{V}_c + 1\right) \cdot \left(1 - \overline{V}^e\right) \cdot \overline{N}} \tag{8}$$

With,

$$\overline{V} = \frac{V}{V^*}, \overline{P} = \frac{P}{P^*}, \overline{N} = \frac{N}{N^*}, \overline{V}_c = \frac{V_c}{V^*}, \overline{\Pi}_0 = \frac{\Delta P + \Pi_0}{E} \tag{9}$$

And,

$$\overline{V}^e = 2\overline{\delta} \cdot (1 + 2\overline{\delta}) \cdot \overline{V}_w + \frac{\overline{R}^*}{\alpha}, \alpha = \frac{\pi/4}{\left(1 + 2 \cdot \overline{\delta}\right)}, \overline{R}^* = \frac{v_p \cdot P^*}{V^*}, \overline{V}_w = \frac{n^* \cdot w^3}{V^*}, \overline{\delta} = \frac{\delta}{w} \tag{10}$$

For convenience, all quantities have been normalized by their values at the time where the confluency constraint is achieved: Eq.3. We highlight that the time of confluency does not correspond to the time at which the pressure starts increasing (Sec.II and Fig.1.C and F). Moreover, we have used that $\overline{P} = \overline{n}$ (see Sec.I D for a justification). Note that Eq.8 was obtained using the following relation:

$$\overline{V}_c \cdot \frac{kTN^*}{\alpha E V^*} = \left((\overline{\Pi}_0 - 1) \cdot \overline{V}_c + 1\right) \cdot \left(1 - \overline{V}^e\right) \tag{11}$$

We model in the next subsections the production of proteins $P$ and osmolytes $N$ to describe how $\overline{V}$ and thus the mechanical stress in the chamber, also called growth induce pressure in the main text (Eq.7), varies with time.

##### B. Proteins production rate

Proteins have been proposed to play a minor role in the osmotic pressure due to their small concentrations in cells relative to small osmolytes (e.g. metabolites) [1]. They nevertheless play an important role in cellular growth by producing/importing those small osmolytes. We thus describe the total protein production rate within the chamber with the following equation:

$$\frac{\mathrm{d}\overline{P}}{\mathrm{dt}} = k_p \cdot \overline{P} \tag{12}$$

Where $k_p$ accounts for the protein growth rate. In the regime where $k_p$ is constant, Eq.12 describes an exponential increase of the proteins, which molecularly arise from the fact that ribosomes autocatalyze their formation. In agreement with previous studies in yeast ([6]), we observe that protein production is sensitive to macromolecular crowding. To include this feature in our growth model and supported by this recent literature, we take that the protein production rate is diffusion-limited and use a Doolittle-like equation ([6]) to relate the variations of the protein production rates with the protein concentration, and thus the macromolecular crowding.

$$k_p = k_p^* \cdot e^{-\xi_p \cdot v_p \cdot (p^b - p^{b,*})} \tag{13}$$

Where, $k_p^*$ is the protein growth rate in the absence of confinement, $v_p$ the average volume of one crowder, $p^b$ and $p^{b,*}$ the protein concentrations respectively during growth and at the onset of bacterial confluency, and $\xi_p$ an adimensional coefficient that describes the sensitivity of protein production to crowding. Using Eq.8, we further express the average bacterial protein concentration as:

$$\frac{p^b}{p^{b,*}} = \frac{\overline{P} \cdot (1 - \overline{V}^e)}{\overline{V} - \overline{V}^e \cdot \overline{P}} \tag{14}$$

##### C. Small osmolytes production rate

In our model, we take into account the production or import of small osmolytes by proteins through the following equation:

$$\frac{\mathrm{d}\overline{\mathrm{N}}}{\mathrm{d}t} = \overline{k}_{osm} \cdot k_p^* \cdot \overline{P} \qquad \text{With,} \quad \overline{k}_{osm} = \frac{k_{osm}}{k_p^*} \cdot \frac{P^*}{N^*} \tag{15}$$

With, $k_{osm}$ the small osmolyte growth rate. We considered two scenarios for the dependency of $k_{osm}$ with crowding. (1) $k_{osm}$ is constant throughout the experiment. (2) Or it decreases exponentially with crowding, identically to the protein growth rate dependency: $k_{osm} = k_{osm}^* \cdot e^{-\xi_p \cdot v_p \cdot (p^b - p^{b,*})}$. Unless specified, in the following we will refer to scenario (1) as it better fits our data (see Supplementary Fig.5).

Note that if proteins grow exponentially, the latter equation predicts that the number of those small osmolytes ruling cell volume, also grows exponentially at the same rate which was recently proposed to explain the origin of protein concentration homeostasis in free unicellular growth ([1]). On the other hand, in the regime of protein production arrest due to crowding, Eq.15 predicts that the number of small osmolytes keeps increasing in the chamber. As the chamber prevents volume growth, this increase in number is further accompanied by an increase in osmotic pressure (Eq.2) which also increases the mechanical stress in the chamber to sustain the pressure balance (Eq.1). This natural decoupling between volume growth and osmolyte production is what we propose to be at the core of the singular regime of growth that we observe with protein production arrest and chamber pressure increase.

##### D. Bacterial number

Important mechanistic details have been understood in recent years on the main drivers leading to bacterial division. In Yeasts and Eukaryotes, a body of evidence shows that the division signal is achieved by the dilution of specific division inhibitors during growth (Whi5 in Yeast, and its mammalian cell homolog, Rb ([7] and [8]). In bacteria, however, the prevailing mechanism is not related to protein concentration but instead to the number of proteins necessary to assemble the cytoskeletal scaffold of the Z ring that constricts to divide the cell in two (see Ftsz protein and [9]). Based on this mechanism and for the sake of simplicity in our model, we propose to implement bacterial division every time the protein number has been doubled. This choice is twofold. (1) It allows for homeostasis of protein concentration during free growth based on a criterion relying on a number of proteins. (2) Most proteins were shown to scale with each other during growth. Under this assumption, the total number of proteins is a proxy of the number of enzymes ruling the number of small osmolytes. Our model would thus remain valid up to a scaling factor that would renormalize the osmolyte production rate, $\overline{k}_{osm}$. According to the previous reasoning, on average, after j divisions the total bacterial protein content has been multiplied by $2^j$ as well as the total number of bacteria:

$$2^j = \overline{P}(t^{d,j}) = \overline{n} \tag{16}$$

Where, $\overline{n}$ accounts for the number of bacteria normalized by the number of bacteria at confluency, and $t^{d,j}$ the time at which the $j^{th}$ division occurs. In the continuum limit, assuming that there are many bacterial divisions at the timescale of our experiments ($\sim 13h$), the previous equation simply states that the number of bacteria is a proxy of the normalized number of proteins that have been produced $\overline{n}(t) = \overline{P}(t)$.

##### E. Summary of the equations

The problem to solve reads:

$$\frac{\mathrm{d}\overline{P}}{\mathrm{d}t} = k_p(\overline{P}, \overline{N}) \cdot \overline{P} \tag{17}$$

$$\frac{\mathrm{d}\overline{N}}{\mathrm{d}t} = \overline{k}_{osm} \cdot k_p^* \cdot \overline{P} \tag{18}$$

With: $\overline{P}(0) = 1$, $\overline{N}(0) = 1$

The other observables can be derived from the solutions of Eq.17 and Eq.18 according to Eq.**??**, Eq.4 and to the following relationships:

$$\sigma^{el} = E \cdot \left( \frac{\overline{V}}{\overline{V}_c} - 1 \right) \quad , \quad n^{2D} = n^{2D,*} \cdot \frac{\overline{n}}{\overline{V}} \tag{19}$$

And,

$$g_d = \frac{1}{\overline{n}_b} \cdot \frac{\mathrm{d}\overline{n}_b}{\mathrm{d}t} = \frac{ln(2)}{\tau_{div}} \quad , \quad g_a = \frac{A_c}{\frac{\pi}{4} \cdot n^{2D,*} \cdot A^b(t=0)} \cdot \frac{1}{\overline{n}} \frac{\mathrm{d}\overline{V}}{\mathrm{d}t} \tag{20}$$

The previous system of equations is solved using the Mathematica software.

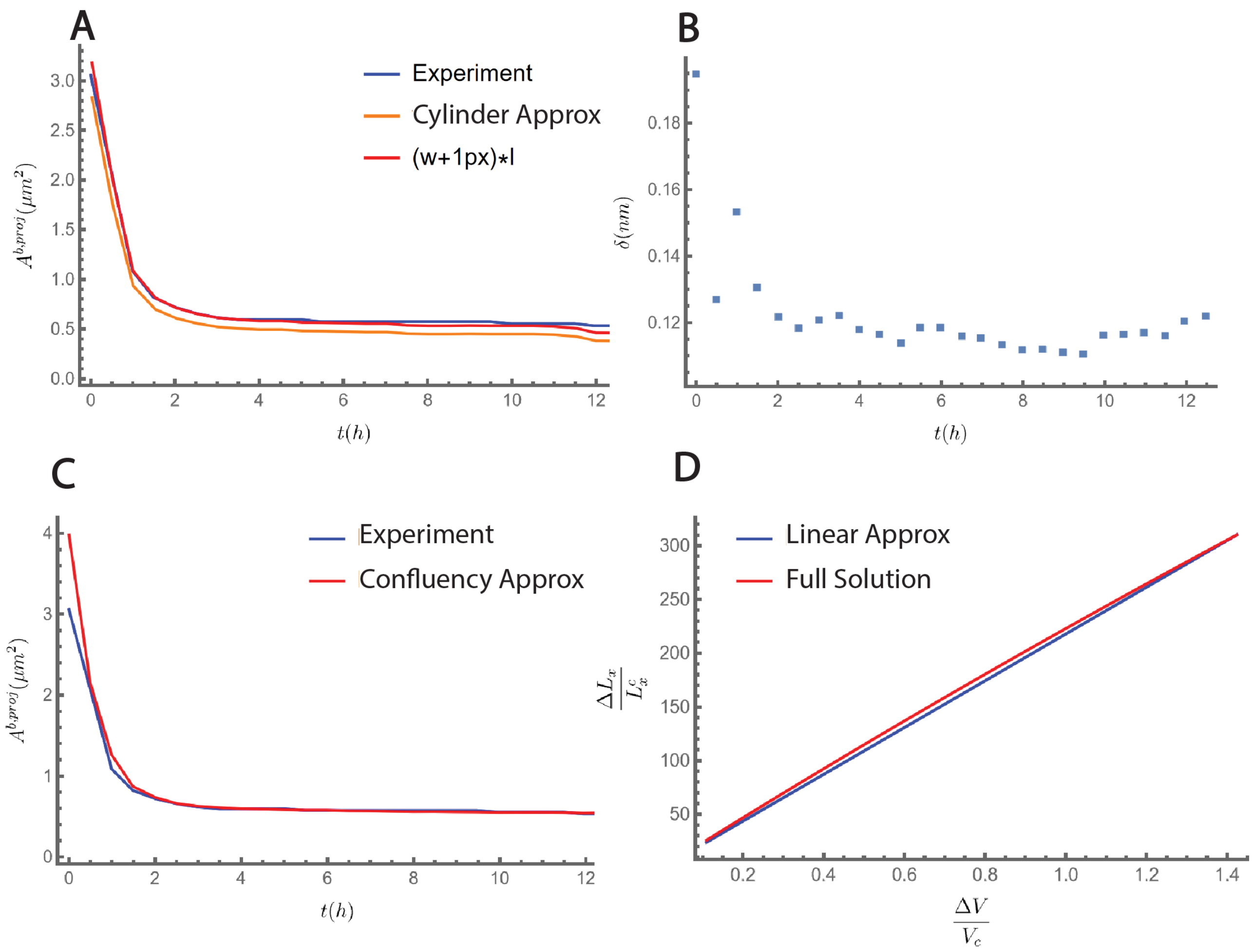


FIG. 1. **Calibration of the model** (A) Comparison between the experimental bacterial projected area (blue) and the projected area assuming that the average shape of the bacteria is a cylinder (orange). This shape approximation is within the experimental measurement error of the actual data as adding one pixel to the experimental width makes the shape approximation match the experimental data. (B) Estimation of the width of the interstitial space using Eq.27 (see Fig.5.B of the main text for a definition). (C) Comparison between the experimental bacterial projected area (blue) and the projected area assuming confluency (Eq.3). Both curves match after 30 minutes. (D) Horizontal versus volume strains under no approximations (red) and following a linear approximation (blue).

#### II. CALIBRATION OF THE MODEL

The model we have developed displays a significant number of parameters.

$$\{t^*, \xi_p, k_p^*, n^{2D,*}, E, \overline{R}^*, \overline{\Pi}_0, \overline{k}_{osm}, \overline{\delta}, \overline{V}_w, \overline{V}_c\} \tag{21}$$

The object of this section is to determine the value of these parameters to decrease the number of adjustable parameters.

##### A. Young modulus of the chamber E

We first calibrate the young modulus of the chamber by applying a known pressure and by measuring the horizontal displacement:

$$\sigma^{el} = E^{eff} \cdot \frac{\Delta L_x}{L_x^c} \tag{22}$$

We find, $E^{eff} = 4.439$MPa. Note that $E^{eff}$ is not equal to the young modulus $E$ defined in Eq. 7 as the horizontal strain of the chamber is not equal to the volume strain. To relate the latter strains we consider the chamber as a truncated pyramid whose upper surface expands with time and thus express the volume of the chamber as:

$$V(t) = h(t) \cdot \frac{\left(Ac + A(t) + \sqrt{Ac \cdot A(t)}\right)}{3} \tag{23}$$

In our experiment the bottom surface cannot move and the surface $A_c$ thus remains equal to its equilibrium value. On the other hand, the upper surface and the height of the chamber expand due to the stress exerted by the bacterial colony growth:

$$A(t) = (L_x^c + \Delta L_x) \cdot (L_y^c + \Delta L_y) \quad And, \quad h(t) = h_c + \Delta h \tag{24}$$

In practice we observe that after 13 hours of growth, $\Delta L_x \sim \Delta L_y \sim \Delta h \sim 2\mu m$. For the sake of precision, we will assume that $\Delta L_y = \alpha_y \cdot \Delta L_x$ and $\Delta L_z = \alpha_z \cdot \Delta L_x$. We measure $\alpha_y \approx 0.93$ and $\alpha_z \approx 1.25$ by making the ratio of the final displacements. By expanding Eq.23 at linear order in the horizontal strain, we show that:

$$\frac{\Delta L_x}{L_x^c} \approx \frac{\frac{\Delta V}{V^c}}{\frac{1}{2} + \frac{1}{2} \cdot \alpha_y \cdot \frac{L_x^c}{L_y^c} + \alpha_z \cdot \frac{L_x^c}{L_z^c}} \tag{25}$$

In practice the previous expansion is a good approximation because we find experimentally that $\frac{\Delta L_x}{L_x^c} < 0.07$. Nevertheless, because the vertical strains in our experiments are high - the chamber height is roughly doubled - we quantify the accuracy of our approximation. We find that the effect of Non-linear terms in the expansion are of order 8% and for completeness we effectively incorporate these non linear terms by multiplying $\frac{\Delta L_x}{L_x^c}$ with a factor $\alpha_{NL}$ that we calibrate to be equal to 0.92. Figure 1.F shows that our linear approximate relationship between horizontal and volume strains is in good agreement with the true relationship. Eq.25 finally allows us to relate the effective young modulus that we measure and the one that we use in our model:

$$E = \alpha_{NL} \cdot \frac{E^{eff}}{\frac{1}{2} + \frac{1}{2} \cdot \alpha_y \cdot \frac{L_x^c}{L_y^c} + \alpha_z \cdot \frac{L_x^c}{L_z^c}} \tag{26}$$

Given the non-deformed dimensions of the chamber $L_x^c = 31.4\,\mu\text{m}, \text{L}_\text{y}^\text{c} = 19.2\,\mu\text{m}, \text{L}_\text{z}^\text{c} = 2.3\,\mu\text{m}$, we find that $E = 218kPa$ (see Fig.1.D).

##### B. Bacteria diameter $w$

Using our tracking algorithm, we are able to measure w at the single bacteria level. In average, we find that $w$ is independent of time once confluency is reached (i.e. maximum of surface occupancy Fig.1.C). In average $w \approx 0.33nm$ (Fig.1.B).

##### C. Interstitial space between bacteria $\delta$

In our experiments, we observe that even after confluency is reached the bacterial protoplasms occupy about 50% of the observed chamber area (see Fig.1.C). We thus take into account this observation in our model by considering a shell around each bacteria of width $\delta$ (see Fig.5.A of the main text). Using Eq.4 we express $\delta$ as:

$$\delta = -\frac{1}{4} \cdot \left(1 + \frac{A^{b,proj}}{w^2}\right) + \sqrt{\frac{1}{16} \cdot \left(1 + \frac{A^{b,proj}}{w^2}\right)^2 + \frac{1}{4w^2} \cdot \left(\frac{Ac}{n^{2D}} - A^{b,proj}\right)} \tag{27}$$

Interestingly, using our measurements of average bacterial number $n^{2D}$, diameter $w$, and protoplasm projected area $A^{b,proj}$ we find that $\delta$ is independent of time once confluency is reached (Fig.1.B). Its median value is $\delta = 117$nm. Note that cell wall width in E.Coli is typically about 80nm. Given that the precision of our measurement is about the size of one pixel $\sim 60nm$, we conclude that most of the interstitial space that we measure is occupied by the cell wall.

##### D. Confluency time $t^*$

In our model we define the time at confluency as the time at which the constraint Eq.3 is reached. We determine $t^*$ by comparing the protoplasm bacterial projected area that we measure thanks to our tracking algorithm and the one estimated using the constraint Eq.3. Before confluency, the latter estimation should overestimate $A^b$ and at confluency both values should match. In Fig.1.E we observe that the confluency time is reached at $t^* \approx 0.51h$.

##### E. Values of the input parameters

We summarize in Table.I the values of the calibrated input parameters of the model.

TABLE I. Description and values of the optimal dimensional parameters of our model.

| Symbol | Typical Value | Meaning |
|---|---|---|
| $n^{2D,*}$ | 158 | Apparent bacteria number in the plan of observation |
| $t^*$ | 0.51h | Time of confluency |
| $k_p^*$ | 3h$^{-1}$ | Bacterial division rate at confluency |
| $\overline{\delta}$ | 0.39 | $\overline{\delta} = \frac{\delta}{w}$ |
| $\overline{V}_c$ | 0.90 | $\overline{V}_c = \frac{V_c}{V_b}$ |
| $\overline{V}_w$ | 0.029 | $\overline{V}_w = \frac{n^{2D,*} \cdot w^2}{A_c}$ |
| $E$ | 218kPa | Elastic modulus of the chamber as defined in Eq.7 |
| $\overline{\Pi}_0$ | 4.42 | $\overline{\Pi}_0 = \frac{\Pi_0}{E}$. With, $\Pi_0 = 963$kPa corresponding to a medium of 390mOsm. |

###### 1. $k_{osm}$ is constant

The remaining three parameters - i.e., $\overline{k}_{osm}$, $\overline{R}^*$ and $\xi_p$ - will be used as adjustable parameters to fit our data. After optimization, we find the optimal values of the adjustable parameters to be:

$$\{\xi_p = 5.7 \quad , \quad \overline{R}^* = 5.3 \cdot 10^{-2} \quad , \quad \overline{k}_{osm} = 3.7 \cdot 10^{-3}\} \tag{28}$$

The previous adimensional parameters can be used to estimate important biological dimensional parameters such as the protein or the small osmolyte concentration. Note that to estimate the protein concentration and number we had to take a protein radius of $r_p \sim 3.5nm$ based on data in the literature [10]. Changing $r_p$ and thus $v_p$ is not critical for the following qualitative conclusion. The values found are summarized in Table.III.

TABLE II. Description and values of the fitted dimensional parameters

| Symbol | Typical Value | Meaning |
|---|---|---|
| $p^*$ | $\sim$ 1.3mMol | protein concentration at confluency |
| $n_{osm}^*$ | $\sim$ 400mMol | osmolyte concentration at confluency |
| $\%_{vol}^{drymass}$ | $\sim$ 12% | Volume fraction of the dry mass, $\%_{vol}^{drymass} = \frac{R^{b,*}}{V^{b,*}}$ |
| $\frac{N^{b,*}}{P^{b,*}}$ | $\sim$ 320 | number fraction of osmolytes other proteins |
| $\frac{k_{osm}}{k_p^*}$ | 1.2 | Ratio of osmolyte over protein production rates |

##### 2. $k_{osm}$ decreases exponentially with crowding

In the other scenario where $k_{osm} = k^*_{osm} \cdot e^{-\xi_p \cdot v_p \cdot (p^b - p^{b,*})}$, the best adjustable parameters we find are:

$$\{\xi_p = 3.2 \quad , \quad \overline{R}^* = 7.0 \cdot 10^{-2} \quad , \quad \overline{k}^*_{osm} = 1.0 \cdot 10^{-2}\} \tag{29}$$

In dimensional unit, we find:

TABLE III. Description and values of the fitted dimensional parameters

| Symbol | Typical Value | Meaning |
|---|---|---|
| $p^*$ | $\sim 1.7$mMol | protein concentration at confluency |
| $n^*_{osm}$ | $\sim 400$mMol | osmolyte concentration at confluency |
| $\%^{drymass}_{vol}$ | $\sim 16\%$ | Volume fraction of the dry mass, $\%^{drymass}_{vol} = \frac{R^{b,*}}{V^{b,*}}$ |
| $\frac{N^{b,*}}{P^{b,*}}$ | $\sim 230$ | number fraction of osmolytes other proteins |
| $\frac{k_{osm}}{k^*_p}$ | 2.3 | Ratio of osmolyte over protein production rates |

#### III. MODEL'S LIMITATIONS

While our model allows to reproduce quantitatively our data, it is worth discussing the discrepancies that we observe for the GIP and the division rate $g_D$. First, the GIP never saturates at long timescales. This is due to the fact that $k_{osm}$ is taken to be constant in our model. This implies that even in the absence of protein production, the chamber would continue to pressiorize because small osmolytes are produced. The modelling of the dependencies of the osmolytes production rate with crowding would however require additional data for calibration. Moreover, it is possible that other effects may take a role in the saturation. For instance, the division rule that we implemented in our model (Sec.I D) is likely to be disrupted at high NC ration due to chromatin sterically preventing the division. We let these hypotheses open for further modelling and future studies.

Second, we note that the division rate in our model decreases faster that the one measured at the single bacterial level in the plan of observation. We have two interpretations for this discrepancy. (1) The theoretical division rate is averaged over the whole chamber, not only on the bacteria in the plan of observation. Given that the bacteria that we observe are closer to the nutrient arrival, we hypothetized that those bacteria are dividing much faster than the one positioned in top layers. This hypothesis of heterogeneity in division rates is supported by observations. We indeed observe that even in the plan of observation, bacteria closer to the edges of the chamber also divides much faster. (2) The measured division rate is obtained by averaging the inverse of the division times of single bacteria in the plan of observation. Importantly, bacteria that have stopped dividing are not considered in this averaging which is likely to biase the average value to larger values compared to the model division rate which considers total population averages.

[1] Romain Rollin, Jean-François Joanny, and Pierre Sens. Physical basis of the cell size scaling laws. *eLife*, 12:e82490, May 2023. Publisher: eLife Sciences Publications, Ltd.
[2] Joël Lemière, Paula Real-Calderon, Liam J Holt, Thomas G Fai, and Fred Chang. Control of nuclear size by osmotic forces in *Schizosaccharomyces pombe*. *eLife*, 11:e76075, jul 2022.
[3] Dan Deviri and Samuel A. Safran. Balance of osmotic pressures determines the nuclear-to-cytoplasmic volume ratio of the cell. *Proceedings of the National Academy of Sciences*, 119(21):e2118301119, May 2022. Publisher: Proceedings of the National Academy of Sciences.
[4] Larisa Venkova, Amit Singh Vishen, Sergio Lembo, Nishit Srivastava, Baptiste Duchamp, Artur Ruppel, Alice Williart, Stephane Vassilopoulos, Alexandre Deslys, Juan-Manuel Garcia Arcos, Alba Diz-Munoz, Martial Balland, Jean-Francois Joanny, Damien Cuvelier, Pierre Sens, and Matthieu Piel. A mechano-osmotic feedback couples cell volume to the rate of cell deformation. *eLife*, 11:e72381, April 2022.
[5] Otger Campas and L. Mahadevan. Shape and Dynamics of Tip-Growing Cells. *Current Biology*, 19(24):2102–2107, December 2009.
[6] Baptiste Alric, Cecile Formosa-Dague, Etienne Dague, Liam J. Holt, and Morgan Delarue. Macromolecular crowding limits growth under pressure. *Nature physics*, 18(4):411–416, April 2022.
[7] Kurt M. Schmoller, J. J. Turner, M. Koivomagi, and Jan M. Skotheim. Dilution of the cell cycle inhibitor Whi5 controls budding-yeast cell size. *Nature*, 526(7572):268–272, October 2015.
[8] Evgeny Zatulovskiy, Shuyuan Zhang, Daniel F. Berenson, Benjamin R. Topacio, and Jan M. Skotheim. Cell growth dilutes the cell cycle inhibitor Rb to trigger cell division. *Science*, 369(6502):466–471, July 2020.
[9] Fangwei Si, Guillaume Le Treut, John T. Sauls, Stephen Vadia, Petra Anne Levin, and Suckjoon Jun. Mechanistic Origin of Cell-Size Control and Homeostasis in Bacteria. *Current Biology*, 29(11):1760–1770.e7, June 2019.
[10] Ron Milo Rob Philipps. *Cell Biology by the numbers.* 2015.

### Supplementary Figures and Legends

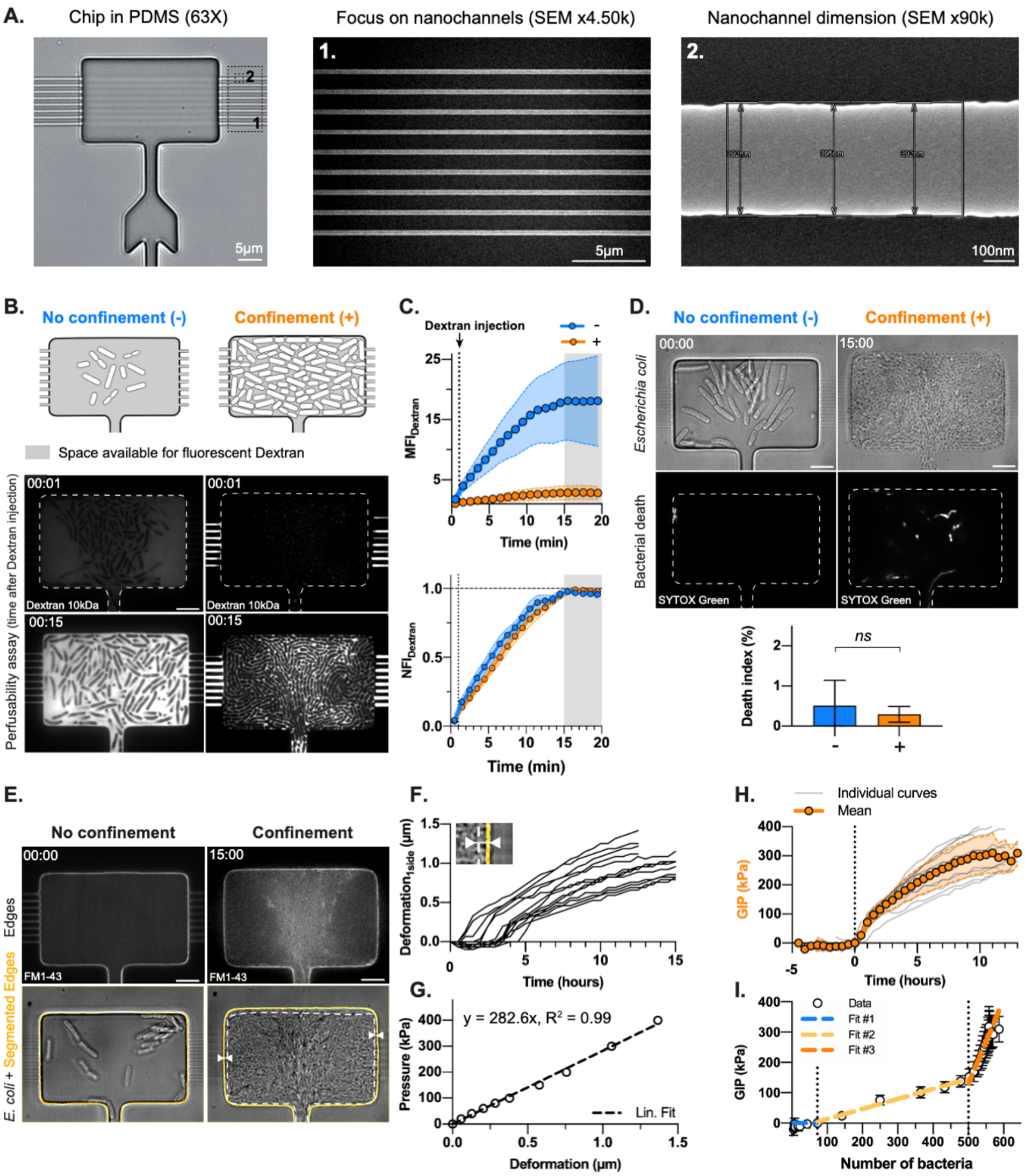


**Supplementary Figure 1**

**Supplementary Figure 1:**

A. Brightfield image of the bacterial confiner by optical microscopy using a 63X objective (*left*). Two regions of interest are highlighted by black dotted squares and corresponding zoom-in are shown on the side. Inset 1. shows a scanning electron microscopy image of the array of 400nm-wide nanochannels connected to the growth chamber (*middle*). Inset 2. shows a scanning electron microscopy image of one single nanochannel for characterization of its dimensions (*right*). B. Schematics of the characterization of aggregate perfusability in the bacterial confiner using 10 kDa Dextran-FITC in absence of confinement (*left*, indicated in blue) and presence of confinement (*right*, indicated in orange). Timelapse confocal images acquired at 1 minute frame rate of Dextran perfusion at different time points of bacterial proliferation in the confiner, before and 3.5h after pressure build-up (*bottom*). Time 0 corresponds to the time 1 min before Dextran injection. (see also **Supp. Video 1**). C. Temporal evolution of Dextran fluorescence inside the growth chamber upon injection, in terms of mean (MFI, *top*) and normalized (NFI, *bottom*) fluorescence intensity. This is calculated in the absence (in blue, $n_{chamber} = 3$, N = 1) or presence of confinement (in orange, $n_{chamber} = 2$, N = 1). The normalized fluorescent intensity is defined as the mean fluorescent intensity divided by its maximum value reached in the time period of 20 minutes after Dextran injection. Time of Dextran injection is depicted by a vertical dotted line. D. Characterization of bacterial viability in the bacterial confiner using the DNA intercalant SYTOX Green dye. Brightfield and confocal images of SYTOX-treated bacteria in the absence and presence of confinement (*top*). Quantification of the death index both in the absence ($n_{FOV} = 48$, $n_{chambers} = 9$, N = 1) and presence of confinement ($n_{FOV} = 199$, $n_{chambers} = 10$, N = 1) (*bottom*). The death index is defined as the percentage of the surface occupied by bacteria which is fluorescently stained by SYTOX Green. Data points correspond to mean values ± standard deviations. Statistical significance of the results was assessed using a Mann-Whitney test. E. Chip optimization to characterize the mechanical environment. Confocal (*top*) and brightfield (*bottom*) images of bacteria growing into a chamber where the walls have been fluorescently stained using the hydrophobic dye FM1-43, in the absence and presence of confinement (0 h and 15 h, corresponding to left and right respectively). Segmented edges of the chamber obtained by semi-automatic detection of the chamber contour over time are highlighted in orange. White arrows and dotted line indicate final chamber deformation respect to its initial size. F. Quantification of the deformation of one wall of the chamber along the x-axis over time, as indicated in the inset. Each curve represents one single chamber (n = 13, N = 3) (see also **Supp. Video 2**). G. Calibration of PDMS deformability, based on the application of a range of known pressures using a pressure controller and measurement of the resulting deformation of a calibration channel along the x-axis. The experimental data points were fitted using a linear regression curve (black dotted line), allowing to relate the measured deformation to a known pressure. H. Quantification of the average growth-induced pressure generated by confined bacteria (n = 13, N = 3). The individual curves (grey) are inferred from the individual deformations in Panel F by using the calibration curve in Panel G and normalized to the time 0 of pressure build-up. Mean values ± standard deviations are represented in orange. I. Relationship between the growth-induced pressure and the number of bacteria measured in the 2D plane of observation. Experimental mean values ± standard deviations are represented. We identified three regimes: in the first one, the number of bacteria increase in the absence of confinement without generating pressure (Fit 1, indicated in blue: $y = 0$). In the second regime, the number of bacteria increases rapidly and the growth-induced pressure also (Fit 2, indicated in yellow: $y = 0.3305\,x - 19.74$, $R^2 = 0.9$). In the third regime, the number of bacteria increases slowly while the pressure continues to increase (Fit 3, indicated in orange: $y = 2.796\,x - 1269$, $R^2 = 0.5$). Time is indicated as hh:mm. Scale bars: 5µm

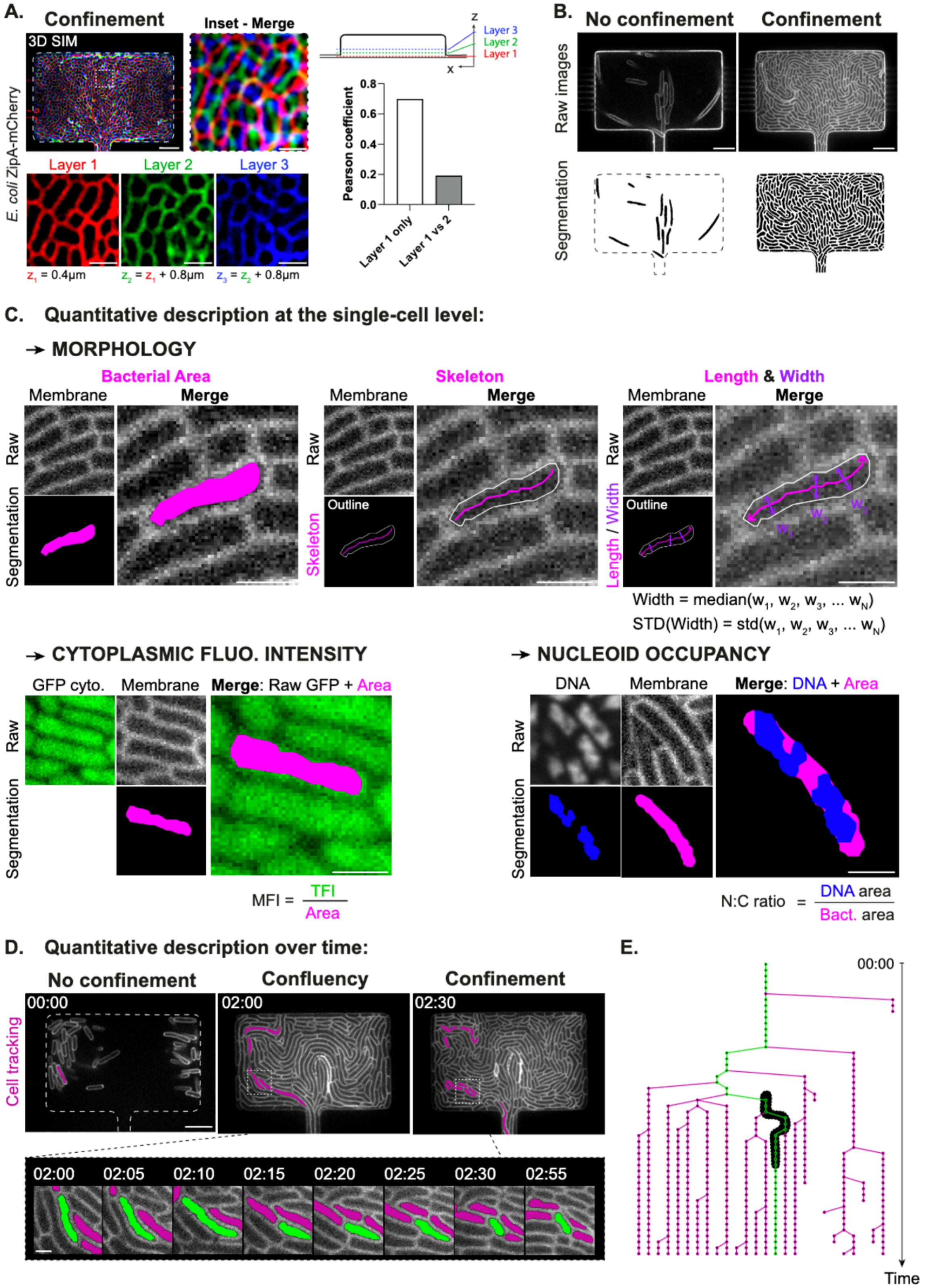


**Supplementary Figure 2**

**Supplementary Figure 2:**
A. 3D SIM super-resolution confocal images of confined *E. coli* MG1655 ZipA-mCherry bacteria fluorescently labeled at the inner membrane in the bacterial confiner after more than 10 h after pressure build-up (*top left*). The inset depicted in white dashed lines focuses on single bacteria (merge: *top middle*, single layers: *bottom left*). The three first layers of bacteria at the bottom of the chamber are indicated in color as indicated on the cross-sectional schematic of the bacterial confiner (*top, right*). The layers are chosen 0.8 µm apart, which corresponds to the bacterial width measured on non-verticalized bacteria while including the membrane signal (width = 0.82 ± 0.06µm, $n_{bacteria} = 10$, N = 1). The Pearson coefficient quantifies the correlation between two images and highlights that bacteria in the focal plane of observation (*i.e.* layer 1) are mostly not verticalized in the bacterial confiner ($n_{chamber} = 1$, N = 1) (*bottom left*). B. Confocal images of the *E. coli* MG1655 ZipA-mCherry bacteria in the bacterial confiner in the absence and presence of confinement (*top left* and *right* respectively). Corresponding segmentation obtained by a semi-automatic machine-learning based pipeline (*bottom*). C. Single-cell quantitative description of bacterial physiology upon confinement. For each parameter, a representative raw image with computed features (magenta) and merge are shown, together with corresponding equations when needed. Parameters include bacterial morphology (On top from left to right: area, skeleton, length and width), cytoplasmic fluorescent intensity and nucleoid occupancy. Scale bars: 5 µm (see also **Supp. Video 3**). D. Quantitative description over time of bacterial physiological changes using single-cell tracking. Timelapse confocal images acquired at 5 minutes frame rate of *E. coli* strain fluorescently labeled at the inner membrane proliferating in the bacterial confiner (*top*). Single-cell tracking by the Fiji plugin TrackMate 7 allows to follow bacterial generations during the course of the experiment, and in particular at the onset of confinement. One bacterial family is depicted in pink. Inset of one specific bacterium depicted in green, undergoing two cycles of division and shortening its length upon pressure build-up (*bottom*). Scale bar: 5µm. Scale bar inset: 1 µm (see also **Supp. Video 3**). E. Lineages corresponding to the bacterial family depicted in pink in Panel D. Each point corresponds to one bacterium at a specific time point. A straight line relates one bacterium at a time $t_1$ to another bacterium at time $t_2$ as the same bacterium. A split in two branches corresponds to bacterial division in two daughter cells. The dark region refers to the time resolved dark inset in Panel D.

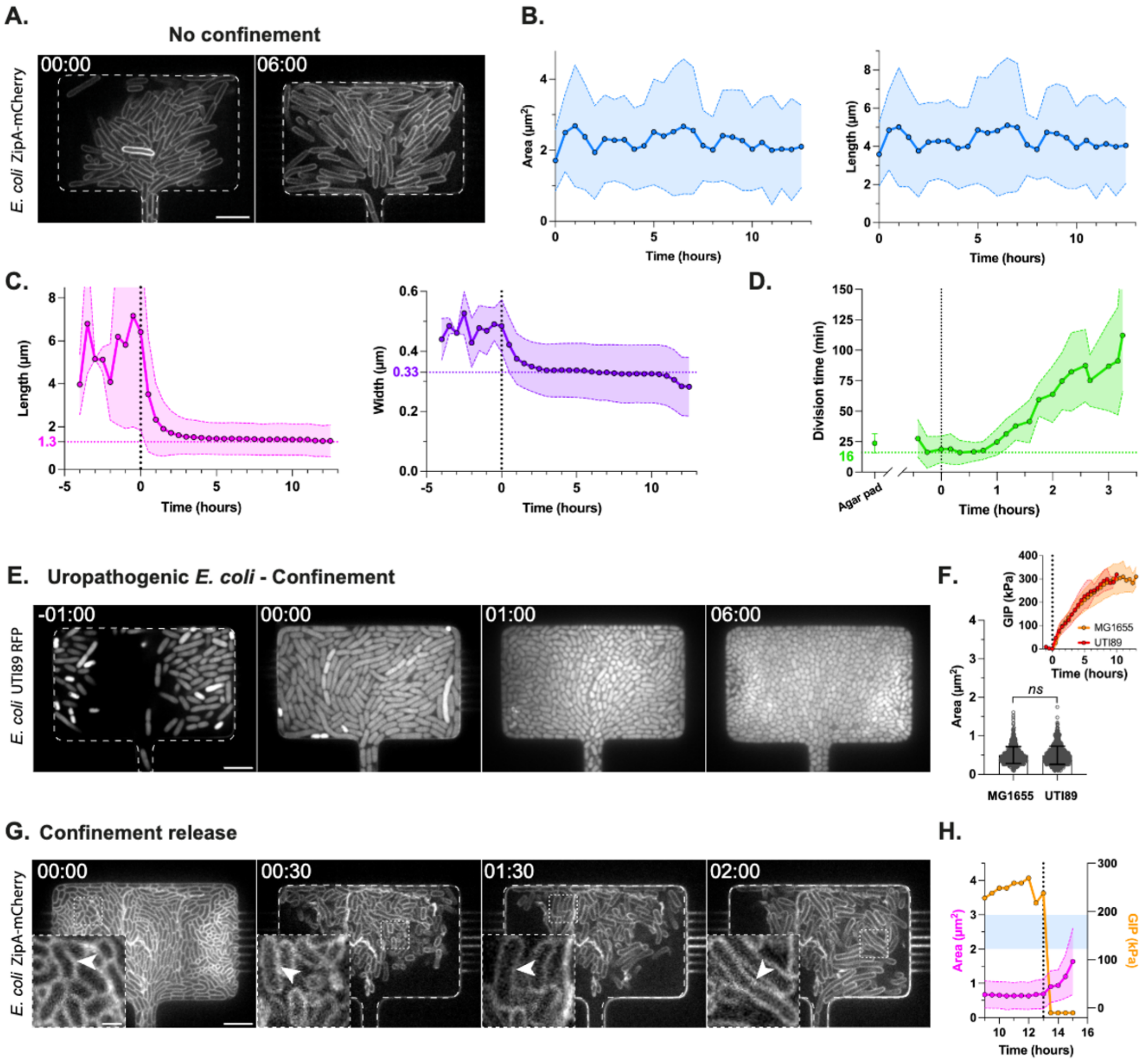


**Supplementary Figure 3**

**Supplementary Figure 3:**
A. Timelapse confocal images acquired at 30 minutes frame rate of the *E. coli* MG1655 strain fluorescently labeled at the inner membrane proliferating in the bacterial confiner in the absence of confinement. B. Quantification of bacterial area (*left*) and length (*right*) in the absence of confinement ($n_{bacteria} = 3871$, $n_{chambers} = 1$, $N = 1$). Mean values ± standard deviations are represented in blue. C. Quantification of bacterial length and width in the presence of confinement ($n_{bacteria} = 47834$, $n_{chambers} = 4$, $N = 2$). Time 0 corresponds to the time of pressure build-up and is indicated by a black dotted line. Mean values ± standard deviations are represented in magenta and purple, and final values reached at late stages of confinement are highlighted. D. Quantification of bacterial division time in agar pad (*left*, $n_{bacteria} = 526$, $n_{tracks} = 68$, $N = 1$) and upon confinement (*right*, $n_{bacteria} = 26728$, $n_{tracks} > 800$, $n_{chambers} = 2$, $N = 1$). Time 0 corresponds to the time of pressure build-up and is indicated by a black dotted line. Mean values ± standard deviations are represented in green, and initial value characterizing the early phase of confinement is highlighted. E. Timelapse confocal images acquired at 30 minutes frame rate of the uropathogenic *E. coli* UTI89 fluorescently labeled in the cytoplasm proliferating in the bacterial confiner in the presence of confinement. Time 0 corresponds to the time of pressure build-up (see also **Supp. Video 4**). F. Quantification of bacterial area upon late confinement (MG1655: t = 13.5 h, UTI89: t = 8 h) for the MG1655 ($n_{bacteria} = 695$, $n_{chamber} = 1$, $N = 1$) and UTI89 strains ($n_{bacteria} = 701$, $n_{chamber} = 1$, $N = 1$). Inset: Quantification of growth-induced pressure generated by the MG1655 (orange) ($n_{chambers} = 13$, $N = 3$) and UTI89 (red) ($n_{chambers} = 5$, $N = 2$) strains. Statistical significance of the results was assessed using Welch ANOVA test. G. Timelapse confocal images acquired at 30 minutes frame rate of the *E. coli* MG1655 strain fluorescently labeled at the inner membrane proliferating in the bacterial confiner upon confinement release. Insets depict bacterial size recovery. H. Quantification of bacterial area (magenta, $n_{bacteria} = 4873$, $n_{chamber} = 1$, $N = 1$) and growth-induced pressure (orange, $n_{chamber} = 1$, $N = 1$) upon confinement release. The black dotted line identifies the time at which confinement is released. The light blue region identifies typical bacterial area in the absence of confinement. Time is indicate as hh:mm. Scale bars: 5 µm. Scale bar inset: 1 µm.

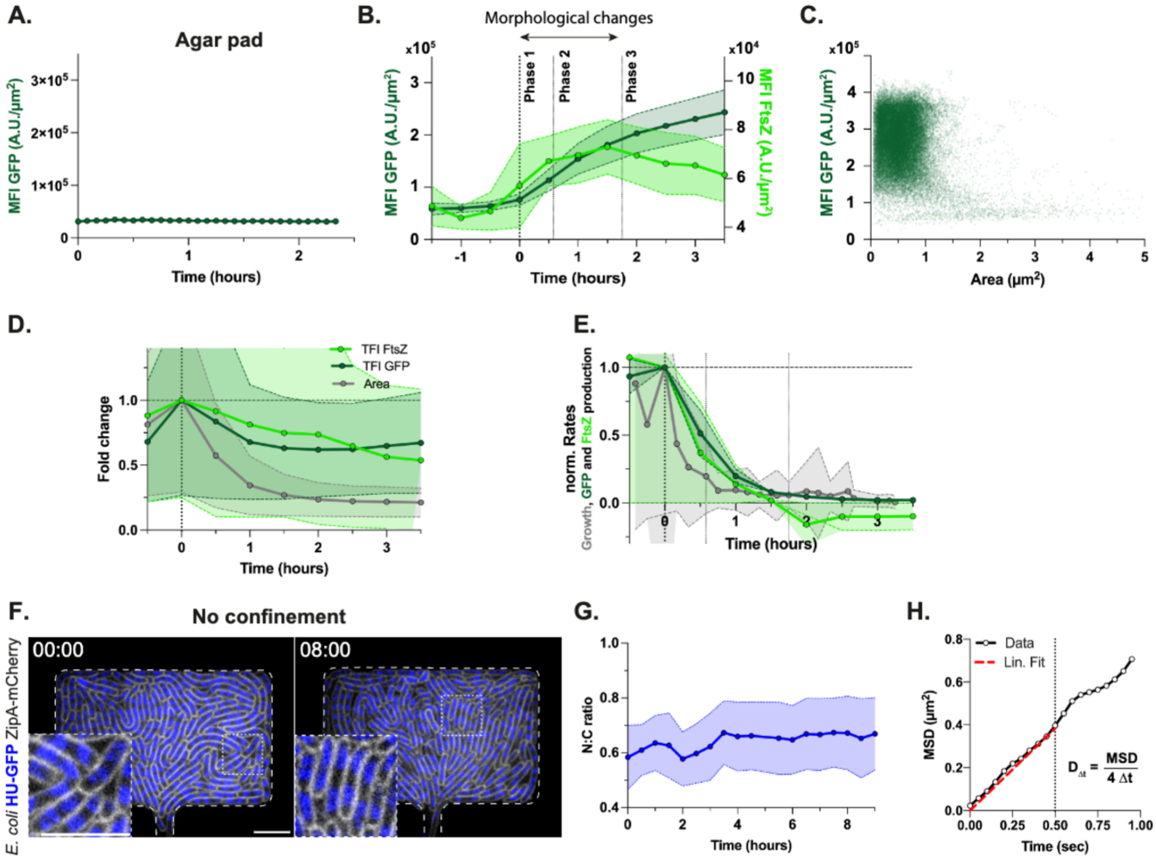


**Supplementary Figure 4**

**Supplementary Figure 4:**
A. Quantification of the GFP mean fluorescent intensity during the proliferation of the *E. coli* MG1655 strain constitutively expressing GFP in agar pad ($n_{bacteria}$ = 1458, N = 1). Mean values ± standard deviations are represented. B. Temporal evolution of single-cell mean fluorescent intensity of the constitutive GFP ($n_{bacteria}$ = 50661, $n_{chambers}$ = 4, N = 1) and FtsZ-mNG ($n_{bacteria}$ = 11855, $n_{chambers}$ = 2, N = 1) signals over the entire cell area respect to pressure build-up, indicated in dark and light green respectively. Phases 1, 2, 3 induced upon confinement are highlighted, together with the time period characterized by major morphological changes. C. Quantification of GFP mean fluorescent intensity as a function of bacterial area upon confinement ($n_{bacteria}$ = 12828, $n_{chamber}$ = 1, N = 1). Each point corresponds to one bacterium at a specific time point. D. Quantification of the fold change in bacterial area, GFP total fluorescent intensity ($n_{bacteria}$ = 50661, $n_{chamber}$ = 4, N = 1) and FtsZ total fluorescent intensity ($n_{bacteria}$ = 11497, $n_{chamber}$ = 2, N = 1) upon confinement. Raw data were normalized by their value at the time of pressure build-up. Mean values ± standard deviations are represented. Bacterial area and FtsZ intensity were measured on different experiments. E. Quantification of total GFP ($n_{chambers}$ = 4, N = 1) and FtsZ ($n_{chambers}$ = 2, N = 1) production rates divided by the bacterial number normalized by their value at time 0 corresponding to pressure build-up (*bottom*). Bacterial growth rate is indicated in grey and computed on another dataset, as shown in Figure 2.D. Data points correspond to mean values ± standard deviations. F. Timelapse confocal fluorescent images acquired at 30 minutes frame rate of the *E. coli* MG1655 HU-GFP ZipA-mCherry strain fluorescently labeled at the inner membrane (grey) and DNA (blue) during proliferation in the bacterial confiner in the absence of confinement. Time is indicated as hh:mm. Scale bar: 5µm. G. Quantification of the karyoplasmic ratio in the absence of confinement. Mean values ± standard deviations are represented in blue ($n_{bacteria}$ = 6025, $n_{chamber}$ = 1, N = 1). H. Quantification of the GEM40 diffusion coefficient. For each individual trajectory, the mean square displacement (written MSD) was plotted as a function of time and fitted with a linear regression curve during the first 0.5 seconds. The slope was then used to infer the corresponding diffusion coefficient based on the equation shown.

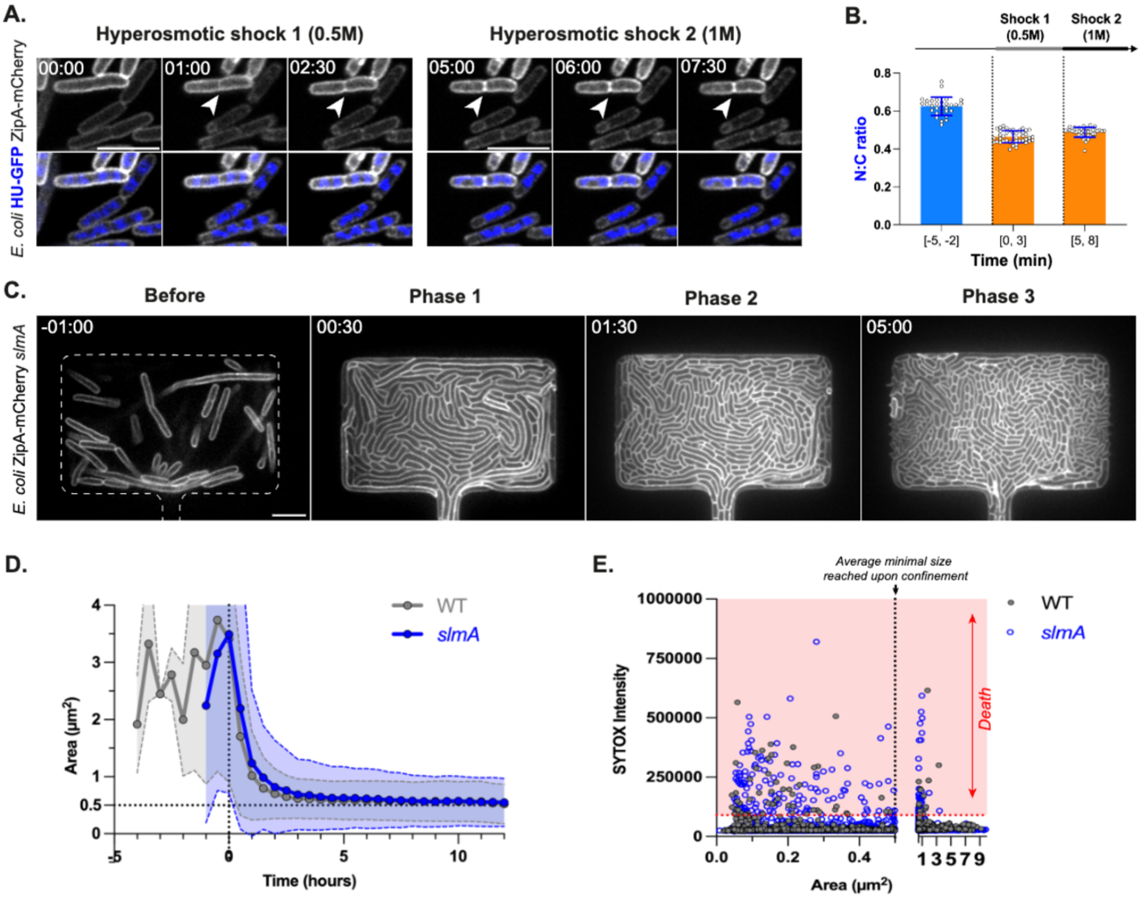


**Supplementary Figure 5**

**Supplementary Figure 5:**

A. Timelapse confocal images acquired at 1 minute frame rate of the *E. coli* MG1655 ZipA-mCherry HU-GFP strain fluorescently labeled at the inner membrane (grey) and DNA (blue) upon two successive hyperosmotic shocks in CellASIC chambers. White arrows indicate the division site. Time is indicated as mm:ss. Scale bars: 5 µm. B. Quantification of the karyoplasmic ratio in response to two successive hyperosmotic shocks (per condition: $n_{bacteria} \geq 199$, $N = 4$). Mean values ± standard deviations are represented. C. Timelapse confocal images acquired at 30 minutes frame rate of the *E. coli* MG1655 ZipA-mCherry *slmA* mutant fluorescently labeled at the inner membrane proliferating in the bacterial confiner. Time is indicated as hh:mm. Scale bars: 5 µm. D. Quantification of bacterial area upon confinement for a wild-type strain (gray) ($n_{bacteria} > 47000$, $n_{chamber} = 4$, $N = 2$) and *slmA* mutant (blue) ($n_{bacteria} = 24960$, $n_{chamber} = 2$, $N = 1$). Mean values ± standard deviations are represented. E. Quantification of SYTOX Green intensity used as a readout of bacterial viability as a function of bacterial area. Each point corresponds to one bacterium at a specific timepoint. The vertical dotted line identifies the average minimal size reached upon late confinement. The region highlighted in red represents the SYTOX Green intensities reached upon bacterial death.

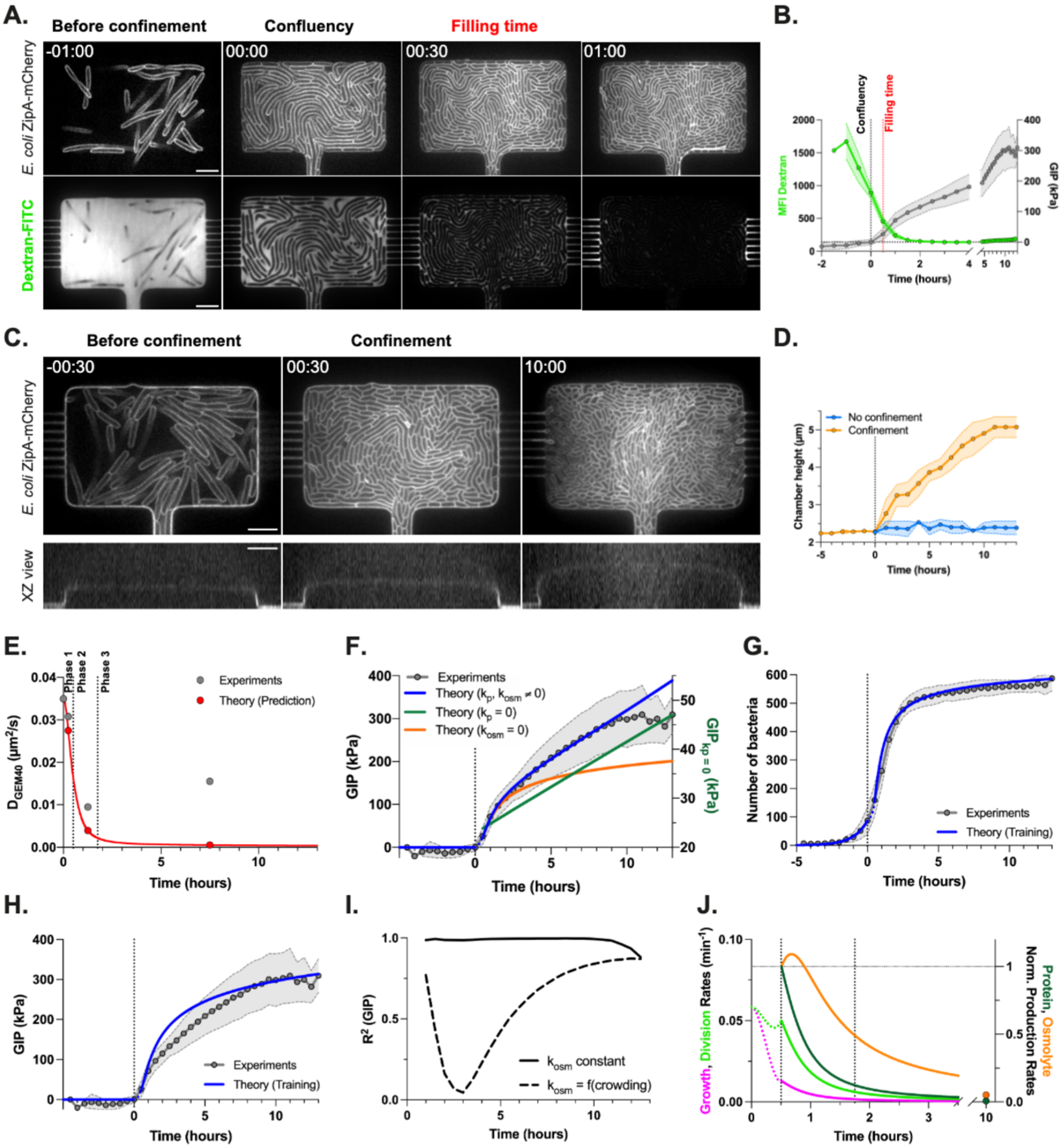


**Supplementary Figure 6**

**Supplementary Figure 6:**

A. Timelapse confocal images acquired at 30 minutes frame rate of *E. coli* MG1655 ZipA-mCherry (*top*) during proliferation in the bacterial confiner in the presence of a culture medium supplemented with Dextran-FITC (*bottom*) to quantify bacterial occupancy. We define confluency as the time at which bacteria become in close contact with their neighbors and start to build up pressure, and filling time as the time at which bacteria fully occupy the 2D bottom plane of the chambers. The filling time is determined as the time at which the signal of the fluorescent medium decreases by 75% compared to its initial value. Note that the filling time takes place 30 minutes after confluency. B. Corresponding temporal evolution of mean fluorescent intensity of Dextran-FITC ($n_{chamber} = 3$, N = 1) and GIP ($n_{chamber} = 13$, N = 3) are indicated in green and grey respectively. Mean values ± standard deviations are represented. C. Timelapse confocal images acquired at 30 min frame rate of *E. coli* MG1655 ZipA-mCherry (*top*) during confined growth and corresponding XZ views of the chamber walls stained with FM1-43 (*bottom*) acquired at 1 h frame rate to quantify deformation in the z direction. D. Corresponding temporal evolution of chamber height in the absence (blue) and presence (orange) of confinement ($n_{chamber} = 3$, N = 1). Mean values ± standard deviations are represented. E. Theoretical prediction (red) and corresponding experimental data points (grey) of GEM40 effective diffusion coefficients upon confinement in Phase 1, 2 and 3. F. Experimental curve of growth-induced pressure (grey) and theoretical fits corresponding to models where $k_p$, $k_{osm} \neq 0$ (blue), $k_p = 0$ (green) and $k_{osm} = 0$ (orange). G. Experimental curve (grey) and theoretical fit (blue) of the number of bacteria in 2D as a function of time, based on a model where osmolyte synthesis ($k_{osm}$) has the same dependance to crowding than protein synthesis, and decreases exponentially with the protein concentration. H. Experimental curve (grey) and theoretical fit (blue) of growth-induced pressure as a function of time, based on a model where osmolyte synthesis ($k_{osm}$) has the same dependance to crowding than protein synthesis, and decreases exponentially with the protein concentration. I. Comparison of fit accuracy in the case of growth-induced pressure for the two models based on $R^2$ ($k_{osm}$ = constant: solid line; $k_{osm}$ = f(crowding): dashed line). J. Theoretical predictions of bacterial growth rate (magenta), division rate (light green), protein normalized production rate (dark green), and osmolyte normalized production rate (orange) in a model where osmolyte synthesis ($k_{osm}$) has the same dependance to crowding than protein synthesis and decreases exponentially with the protein concentration. Note that in this case, protein growth rate decreases faster than the small osmolytes growth rate upon confined growth, supporting the overpressurization of bacterial growth for long periods under protein production arrest. Time hh:mm. Scale bar: 5µm

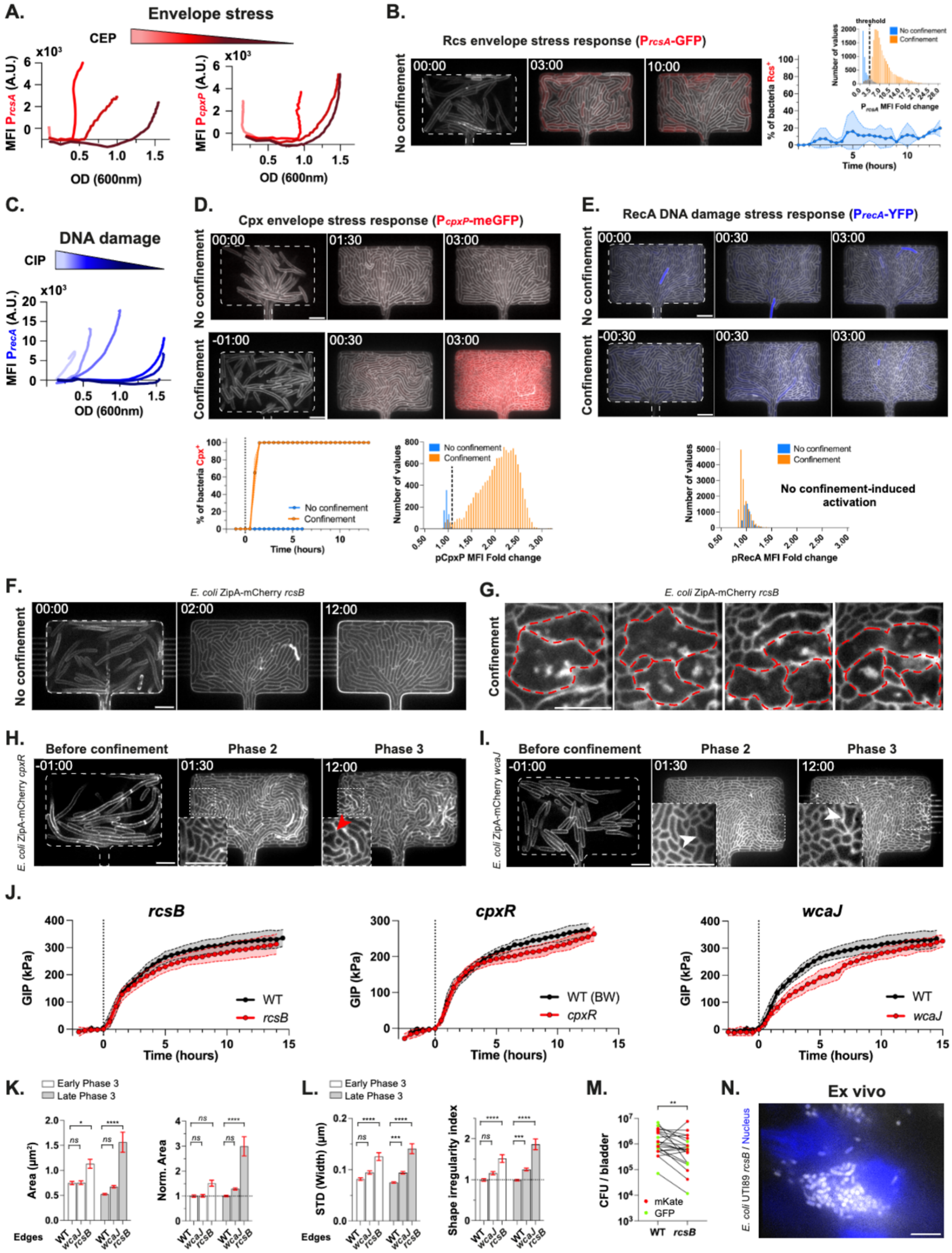


**Supplementary Figure 7**

**Supplementary Figure 7:**

A. Validation of $P_{rcsA}$-GFP (*left*) and $P_{cpxP}$-meGFP (*right*) transcriptional fluorescent reporters upon chemical-induced envelope stress in liquid cultures by using a Cytation plate reader. Quantification of the reporter mean fluorescent intensity as a function of the optical density for a range of cephalexin (CEP) concentrations (0, 5, 15, 20 µg/ml). At a given optical density, the higher the concentration of the chemical inducer, the brighter the fluorescent reporter. B. Timelapse confocal images acquired at 30 minutes frame rate of the *E. coli* MG1655 ZipA-mCherry $P_{rcsA}$-GFP strain fluorescently labeled at the inner membrane (grey) expressing the $P_{rcsA}$ fluorescent reporter (red) while proliferating in the bacterial confiner in the absence of confinement (*left*). Quantification of the percentage of bacteria expressing the Rcs stress response in the absence of confinement ($n_{total\ bacteria}$ = 35200, $n_{chambers}$ = 3, N = 1) (*right*). The number of bacteria activating the Rcs stress response is determined by using a threshold on the mean fluorescent intensity fold change. In the inset, histograms of the $P_{rcsA}$ reporter mean fluorescent intensity fold change for non-confined (blue) and confined bacteria (orange). In the study, a fluorescent reporter is activated as soon as these two histograms form two discernable peaks. In this case, a threshold above which the bacteria activate the stress response is defined and allows to compute the percentage of positive bacteria. C. Validation of $P_{recA}$-YFP transcriptional fluorescent reporter upon chemical-induced DNA damage in response to a range of ciprofloxacin (CIP) concentrations (0, 0.01, 0.1, 0.5, 1, 10 µg/ml) in liquid cultures by using a Cytation plate reader. D. Timelapse confocal images acquired at 30 minutes frame rate of the *E. coli* MG1655 ZipA-mCherry $P_{cpxP}$-meGFP strain fluorescently labeled at the inner membrane (grey) expressing the $P_{cpxP}$-meGFP transcriptional reporter (red) while proliferating in the bacterial confiner in the absence and in the presence of confinement (*top*). Time 0 indicate the time of pressure build-up. Quantification of the percentage of bacteria activating the Cpx stress response in the absence of confinement (blue, $n_{total\ bacteria}$ = 160, $n_{chamber}$ = 1, N = 1) or upon confinement (orange, $n_{total\ bacteria}$ = 41160, $n_{chambers}$ = 3, N = 1) (*bottom*). The number of bacteria expressing the Cpx stress response is determined by using a threshold on the mean fluorescent intensity fold change. In the inset, histograms of the $P_{cpxP}$ reporter mean fluorescent intensity fold change for non-confined and confined bacteria (indicated in blue and orange respectively) used to determine this threshold (see also **Supp. Video 6**). E. Timelapse confocal images acquired at 30 minutes frame rate of the *E. coli* MG1655 ZipA-mCherry $P_{recA}$-YFP strain fluorescently labeled at the inner membrane (grey) expressing the $P_{recA}$ transcriptional fluorescent reporter (blue) during proliferation in the bacterial confiner in the absence and in the presence of confinement (*top*). Time 0 indicate the time of pressure build-up. Histograms of $P_{recA}$ reporter mean fluorescent intensity fold change for non-confined (blue) and confined bacteria (orange) (*bottom*). Since these two histograms fully overlap, we consider that this stress response is not activated upon confinement. (see also **Supp. Video 6**). F. Timelapse confocal images acquired at 30 minutes frame rate of the *E. coli* MG1655 ZipA-mCherry *rcsB* mutant fluorescently labeled at the inner membrane (grey) proliferating in the bacterial confiner in the absence of confinement, where no morphological changes occur. G. Confocal images zooming on representative confined bacteria deficient in Rcs stress response. These bacteria are characterized by blebbing, protruding morphologies depicted in red, which are often characterized by pools of membrane in the bacterial cytoplasm. H. Timelapse confocal images acquired at 30 minutes frame rate of the *E. coli* MG1655 ZipA-mCherry *cpxR* mutant fluorescently labeled at the inner membrane (grey) proliferating in the bacterial confiner in presence of confinement. In the insets, *cpxR* bacteria under confinement exhibit minor shape defects as depicted by a red arrow (see also **Supp. Video 7**). I. Timelapse confocal images acquired at 30 minutes frame rate of the *E. coli* MG1655 ZipA-mCherry *wcaJ* non-capsulated mutant fluorescently labeled at the inner membrane (grey) proliferating in the bacterial confiner in the presence of confinement. Representative examples of the phases before confinement

(1 h), 2 and 3 (1.5 h and 12 h after pressure build-up respectively) are shown. Insets zoom in the regions depicted with a white dashed square line. White arrows indicate bacteria with morphological defects (see also **Supp. Video 7**). J. Quantification of growth-induced pressure build-up for all the mutants used in this study (red), including *rcsB* ($n_{chambers} = 25$, N = 1), *cpxR* ($n_{chambers} = 22$, N = 1) and *wcaJ* ($n_{chambers} = 5$, N = 1), and comparison to the corresponding wild-type strain (black) ($n_{chambers} \geq 13$, N = 1). K. Quantification of bacterial area (*left*) and normalized bacterial area (*right*) for wild-type, *wcaJ* and *rcsB* strains at the edges of the chamber at early phase 3 (2 h after GIP build-up, white) and late phase 3 (10 h after GIP build-up, grey). Bacterial area is used as a readout of bacterial size. L. Quantification of standard deviation of bacterial width (*left*) and corresponding shape irregularity index (*right*) for wild-type, *wcaJ* and *rcsB* strains at the edges of the chamber at early phase 3 (2 h after GIP build-up, white) and late phase 3 (10 h after GIP build-up, grey). Standard deviation of bacterial width is used as a readout of bacterial shape irregularity. Means ± standard errors are represented. These quantitative parameters were measured for the wild-type strain ($n_{bacteria – Early Phase 3} = 158$, $n_{bacteria – Late Phase 3} = 478$, $n_{chambers} = 3$, N = 1), *wcaJ* ($n_{bacteria – Early Phase 3} = 164$, $n_{bacteria – Late Phase 3} = 409$, $n_{chambers} = 3$, N = 2) and *rcsB* mutants ($n_{bacteria – Early Phase 3} = 69$, $n_{bacteria – Late Phase 3} = 134$, $n_{chambers} = 2$, N = 1). Mean values ± standard errors are represented. Statistical significance of the results is assessed using one-way ANOVA tests. M. *In vivo* competition experiment in a mouse model of urinary tract infection. Infections were performed by mixing two pairs of UTI89 bacterial strains, WT and *rcsB*, each of those in the same background, expressing GFP and mKate as indicated in green and red respectively. CFU/bladder of *rcsB* and WT strains after tissue dissection and plating on corresponding selective antibiotics are represented and connected as pairs when they come from the same infected mice. Data obtained from all competition experiments were pooled together for statistical analysis, performed using Wilcoxon matched-pairs rank test (to compare CFU from the same bladder – paired data). N. Representative confocal image of an infected uroepithelium at 24 h post-infection showing a tight aggregate of the uropathogenic *E. coli* UTI89 *rcsB* mutant strain confined within the tissue (grey: bacteria; blue: nuclei). Time is indicated as hh:mm. All scale bars: 5 µm.

#### Table 1: Strains and plasmids used in this study

| Strains or plasmids | Description | Relevant information | Source or reference |
|---|---|---|---|
| ***E. coli*** | | | |
| MG1655 | Wild-type *E. coli* K-12 | | Laboratory collection |
| TB28::attHKpHC503 | MG1655 Δ*lacIZYA* $P_{zapA}$-ZipA(TM)-mCherry2, $Ap^R$ | source of ZipA-mCherry2 | [1] |
| CHN128 = MG1655 GFP | *intC*::$P_R$-meGFP $Cm^R$ | | [2] |
| SJ156 = HU-GFP $P_{sulA}$-mCherry | *hupA*-GFP $Km^R$ $P_{sulA}$-mCherry | source of HU-GFP | [3] |
| MG1655 $P_{cpxP}$-meGFP | *galK*::*cpxP*-meGFP $Cm^R$ | | Ivan Matic's lab |
| MG1655 $P_{recA}$-YFP | *intC*::$P_{recA}$-YFP $Cm^R$ | | Ivan Matic's lab |
| MG1655 mKate | Δ*intC*::PR-mKate-*cat* $Cm^R$ | source for mKate | [2] |
| BW25113 *slmA* | Δ*slmA*::$Km^R$ | source for *slmA* | [4] |
| BW25113 *rcsB* | Δ*rcsB*::$Km^R$ | source for *rcsB* | [4] |
| BW25113 *wcaJ* | Δ*wcaJ*::$Km^R$ | source for *wcaJ* | [4] |
| BW25113 *cpxR* | Δ*cpxR*::$Km^R$ | source for *cpxR* | [4] |
| BW27783 FtsZ-mNG | $P_{ftsZ}$-G55-*ftsZ*-Q56-mNeonGreen | | [5] |
| XL1-Blue | *recA1 endA1 gyrA96 thi-1 hsdR17 supE44 relA1 lac* [F´ *proAB lacI*$^q$ *Z*Δ*M15* Tn*10*] $Tc^R$ | | Stratagene |
| MG1655 ZipA-mCherry | *tet*-$P_{zapA}$-ZipA(TM)-mCherry2 $Tc^R$ | *attB*$_{HK022}$ site | This study |
| MG1655 GFP ZipA-mCherry | *intC*::$P_R$-meGFP $Cm^R$ *tet*-$P_{zapA}$-ZipA(TM)-mCherry2 $Tc^R$ | | This study |
| MG1655 HU-GFP | *hupA*-GFP $Km^R$ | | This study |
| MG1655 HU-GFP ZipA-mCherry | *hupA*-GFP $Km^R$ *tet*-$P_{zapA}$-ZipA(TM)-mCherry2 $Tc^R$ | | This study |
| BW27783 FtsZ-mNG ZipA-mCherry | $P_{ftsZ}$-G55-*ftsZ*-Q56-mNeonGreen *tet*-$P_{zapA}$-ZipA(TM)-mCherry2 $Tc^R$ | | This study |
| MG1655 $P_{cpxP}$-meGFP ZipA-mCherry | *tet*-$P_{zapA}$-ZipA(TM)-mCherry2 $Tc^R$ *galK*::*cpxP*-meGFP $Cm^R$ | | This study |
| MG1655 $P_{recA}$-YFP ZipA-mCherry | *tet*-$P_{zapA}$-ZipA(TM)-mCherry2 $Tc^R$ *intC*::$P_{recA}$-YFP $Cm^R$ | | This study |
| XL1-Blue GEM40 | *cat*-$P_{lac}$-PfV-GS-Sapphire $Cm^R$ | *attB*λ site | This study |
| MG1655 GEM40 | *cat*-$P_{lac}$-PfV-GS-Sapphire $Cm^R$ | *attB*λ site | This study |
| MG1655 GEM40 ZipA-mCherry | *cat*-$P_{lac}$-PfV-GS-Sapphire $Cm^R$ *tet*-$P_{zapA}$-ZipA(TM)-mCherry2 $Tc^R$ | | This study |
| MG1655 ZipA-mCherry *slmA* | *tet*-$P_{zapA}$-ZipA(TM)-mCherry2 $Tc^R$ Δ*slmA*::$Km^R$ | | This study |
| MG1655 ZipA-mCherry *rcsB* | *tet*-$P_{zapA}$-ZipA(TM)-mCherry2 $Tc^R$ Δ*rcsB*::$Km^R$ | | This study |
| MG1655 ZipA-mCherry *wcaJ* | *tet*-$P_{zapA}$-ZipA(TM)-mCherry2 $Tc^R$ Δ*wcaJ*::$Km^R$ | | This study |
| MG1655 ZipA-mCherry *cpxR* | *tet*-$P_{zapA}$-ZipA(TM)-mCherry2 $Tc^R$ Δ*cpxR*::$Km^R$ | | This study |
| in UTI89 (UPEC) | | | |
| UTI89 RFP | *aphA*-marsRFP $Km^R$ | | [6] |
| UTI89 GFP | *bla*-GFP $Ap^R$ | | [6] |
| UTI89 GFP *rcsB* | *bla*-GFP $Ap^R$ Δ*rcsB*::$Km^R$ | | This study |
| UTI89 | | $Str^R$ | [7] |
| UTI89 mKate | Δ*intC*::PR-mKate-*cat* $Cm^R$ | $Str^R$ | This study |
| UTI89 mKate *rcsB* | Δ*intC*::PR-mKate-*cat* $Cm^R$ Δ*rcsB*::$Km^R$ | $Str^R$ | This study |
| **Plasmids** | | | |
| p$P_{rcsA}$-GFP | $P_{rcsA}$-*gfpmut2* | | [8] |
| pKOBEG | pSC101$^{ts}$Ω$P_{BAD}$-*red*$\gamma\beta\alpha$ for λ Red recombination $Cm^R$ | | [9] |
| pKOBEGA | pSC101$^{ts}$Ω$P_{BAD}$-*red*$\gamma\beta\alpha$ for λ Red recombination $Ap^R$ | | [9] |
| pBR322 | ColE1 origin, $Ap^R$ $Tc^R$ | source of *tet* gene | [10] |
| pCDNA3.1-pCMV-PfV-GS-Sapphire | Fusion of the gene encoding for *Pyrococcus furiosus* encapsulin with the Sapphire fluorophore, $Ap^R$ | source of GEM40 Addgene #116933 | [11] |
| pMGC10 | Cloning vector for GEM40, LacI repressor and $P_{lac}$ $Ap^R$ $Cm^R$ | | [12] |
| pREP4 | p15A origin, LacI repressor $Km^R$ | | Qiagen |

#### Table 2: List of the primers used in this study

| Primer name | Sequence (5'→ 3') (enzymes restriction sites are underlined) |
|---|---|
| HK022-att-Tc-F | CCATCCAGAGTCTTCGGGTCAGGGTTAAATTCACGGTCGGTGCACTTTAGTTTGCGCATTCACAGTTCTC |
| Tc-R | AGTGGTAGTCTCATGTTTGACAGCTTATCATCG |
| Tc-PzapA-F | GATAAGCTGTCAAACATGAGACTACCACTTTGGGCCCTGG |
| HK022-att-mCherry-R | ATAAGGCTTTATGCTAGATGCATTCCGCTTTGCGACTCAACCTTTTTCACAGAAAGGCCGGGAAATACC |
| P1 | GGAATCAATGCCTGAGTG |
| P4 | GGCATCAACAGCACATTC |
| GEM40-XhoI-F | CTCGAGAGGAGTAATTTTATGCTCTCAATAAATCCAACC |
| GEM40-PmeI-R | GTTTAAACTTATTTGTACAATTCATCAATACCA |
| GEM40-seq1-F | CCAAAAAGATATGCAAAGTTGCT |
| GEM40-seq2-R | ACCCACTACCGCTGCCAGACCA |
| lambda-att-Cm-F | CTTTTTGTCTTTTTACCTTCCCGTTTCGCTCAAGTTAGTACAAGAATTGCCGGCGGAT |
| Cm-UGB1479-R | TGAGACGTTGATCGGCACGTAAG |
| Cm-UGB1479-F | GCCGATCAACGTCTCAGACATCATAACGGTTCTGGCAAA |
| lambda-att-Sapphire-R | ATGAAATAGAAAAATGAATCCGTTGAAGCCTGCTTTTTTATTATTTGTACAATTCATCAATACCATGG |
| lambda-att-ext5-F | GGCGATAAATTGCCGCATCG |
| lambda-att-ext3-R | TGCCACCATCAAGGGAAAGCCC |
| hupA-F | ATGAACAAGACTCAACTGATTGAT |
| yjaH-R | GAAGAGTTATGACTACAGGCAGTG |
| slmA-F | GGTCACTCTGGTCGTCAGTA |
| slmA-R | TGATTACAACTGCAAAGTGATCT |
| wcaJ-F | ATGGCATCAGCATGACCTTT |
| wcaJ-R | TCAGAAAGCGTATCTGCCAG |
| cpxR-F | TAACTATGCGCATCATTTGC |
| cpxR-R | GACATGCTGCTCAATCATCA |
| mKate-F | ACGCCAAAAAGCAGAACGTGC |
| mKate-R | TGCACTGGATTGCAAGACTTT |
| rcsB-F | CTTAAAGGCGTATTTGCCAT |
| rcsB-R | GATAAAACAGACGCTGACGTTA |

#### Supplementary Videos - Legends

**Supplementary Video 1: Aggregates perfusability in the bacterial confiner**
Timelapse confocal images acquired at 1 minute frame rate of aggregate perfusability in the bacterial confiner using 10 kDa Dextran-FITC in absence of confinement (indicated in blue) and presence of confinement (3.5h after pressure build-up, indicated in orange). Time 0 corresponds to the time 1 min before Dextran injection. Time is indicated as hh:mm. Scale bar: 5µm. See also **Supp. Figure 1.B**.

**Supplementary Video 2: Bacterial proliferation under confinement generates growth-induced pressure (GIP)**
Timelapse brightfield images acquired at 30 minutes frame rate and segmented chamber edges of bacterial proliferation in the bacterial confiner in the absence of confinement (indicated in blue, *left*), and presence of confinement (indicated in orange, *right*). The frame corresponding to the onset of growth-induced pressure (GIP) is highlighted. Time is indicated as hh:mm. Scale bar: 5µm. See also **Figure 1.B-C**.

**Supplementary Video 3: Growth-induced pressure leads to rod cell shortening due to uncoupling between bacterial growth and division**

Timelapse confocal images acquired at 30 minutes frame rate of *E. coli* MG1655 ZipA-mCherry bacteria fluorescently labeled at the inner membrane in the absence of confinement (indicated in blue, *left*), and presence of confinement (indicated in orange, *right*), and corresponding colormap of the bacterial area after single-cell segmentation. Finally, representative inset of a sequence of reductive divisions induced by pressure build-up in the chamber. The frame corresponding to the onset of growth-induced pressure (GIP) is highlighted. Time is indicated as hh:mm. Scale bar: 5µm. See also **Figure 2.A**.

**Supplementary Video 4: Confinement-induced morphological changes are reversible upon pressure release**
Timelapse confocal images acquired at 30 minutes frame rate of pathogenic *E. coli* UTI89 RFP bacteria before, during confinement and after pressure release. The frame corresponding to the onset of growth-induced pressure (GIP) is highlighted. Time is indicated as hh:mm. Scale bar: 5µm. See also **Supp. Figure 4.E**.

**Supplementary Video 5: Confinement increases cytoplasmic crowding through changes in protein concentrations and DNA occupancy**

In a sequence, timelapse confocal images of *E. coli* MG1655 GFP ZipA-mCherry bacteria expressing in their cytoplasm a constitutive GFP (to note, no autoscale/frame was applied in the corresponding video), *E. coli* HU-GFP ZipA-mCherry bacteria fluorescently labeled at the inner membrane and DNA and *E. coli* GEM40 ZipA-mCherry bacteria expressing diffusive nanoparticles in their cytoplasm during proliferation in the bacterial confiner. When required, conditions in the absence of confinement (in blue) and presence of confinement (in orange) are shown. The frame corresponding to the onset of growth-induced pressure (GIP) is highlighted. Time is indicated as hh:mm. Scale bar: 5µm. See also **Figure 3.A, C, E**.

**Supplementary Video 6: Transcriptional stress responses upon mechanical confinement**
In a sequence, timelapse confocal images acquired at 30 minutes frame rate of the *E. coli* MG1655 ZipA-mCherry strain expressing $P_{rcsA}$-GFP, $P_{cpxP}$-meGFP, and $P_{recA}$-YFP during proliferation in the bacterial confiner in the absence and in the presence of confinement (top

left and bottom left respectively). The frame corresponding to the onset of growth-induced pressure (GIP) is highlighted. Time is indicated as hh:mm. Scale bar: 5µm. See also **Figure 6.A**, **Supp. Figure 8.D, E**.

**Supplementary Video 7: Morphology of cell envelope mutants upon mechanical confinement**

In a sequence, timelapse confocal images acquired at 30 minutes frame rate of the *E. coli* MG1655 ZipA-mCherry mutant strains in *rcsB*, *cpxR* and *wcaJ* during proliferation in the bacterial confiner. Conditions in the absence of confinement (indicated in blue, *left*) and presence of confinement (indicated in orange, *right*) are shown. The frame corresponding to the onset of growth-induced pressure (GIP) is highlighted. Time is indicated as hh:mm. Scale bar: 5µm. See also **Figure 6.C**, **Supp. Figure 9.D, E**.
